# An overexpression platform reveals the functional diversity of human KRAB-Zinc Finger Proteins in maintaining cellular homeostasis

**DOI:** 10.64898/2026.04.20.718945

**Authors:** Romain Forey, Charlène Raclot, Arianna Dorschel, Jérôme Archambeau, Evarist Planet, Olimpia Bompadre, Sandra Offner, Wayo Matsushima, F. Gisou van der Goot, Didier Trono

## Abstract

Kruppel associated box zinc finger proteins (KZFPs) form the largest family of transcriptional regulators in mammals, yet most remain uncharacterized. Here we established a scalable framework to probe KZFP function. An arrayed inducible overexpression screen of 366 human KZFPs in K562 cells identified factors that alter cellular proliferation, enabling functional prioritization. Integrative transcriptomic, chromatin and proteomic analyses revealed diverse mechanisms, including transposable element–linked repression (ZNF43), promoter proximal regulation (ZNF257), and SCAN domain dependent transcriptional activation (ZNF498/ZSCAN25 and ZNF18). These results highlight the functional diversity of KZFPs and provide a strategy for their annotation.

## Introduction

Krüppel-associated box (KRAB) zinc finger proteins (KZFP) emerged in the last common ancestor of lung fish, coelacanth and tetrapods, and amplified through gene and segment duplication to constitute the largest family of DNA-binding transcriptional regulators encoded by extant mammals, with ∼380 members in human alone. (Urrutia, 2003; Huntley *et al*., 2006; Nowick and Stubbs, 2010; de Tribolet-Hardy *et al*., 2023). Characterized by an N-terminal KRAB domain and a C-terminal tandem array of C2H2 zinc fingers, KZFPs primarily target transposable element (TE)-embedded sequences (Wolf and Goff, 2009; Jacobs *et al*., 2014; Najafabadi, Albu and Hughes, 2015; Schmitges *et al*., 2016; Imbeault, Helleboid and Trono, 2017; Bruno, Mahgoub and Macfarlan, 2019; Helleboid *et al*., 2019; de Tribolet-Hardy *et al*., 2023). Approximately 80% of them contain a “standard” KRAB domain and recruit TRIM28 (KAP1), which serves as a scaffold for a heterochromatin-inducing complex comprising the histone methyltransferase SETDB1, histone deacetylases, the nucleosome remodelling NuRD and the heterochromatin protein 1 (HP1) (Friedman *et al*., 1996; Ryan *et al*., 1999; Schultz, Friedman and Rauscher, 2001; Schultz *et al*., 2002; Birtle and Ponting, 2006). This results in silencing of DNA elements targeted by these KZFPs, as notably observed for TEs during early embryogenesis (Wolf and Goff, 2009; Matsui *et al*., 2010; Rowe *et al*., 2010; Castro-Diaz *et al*., 2014; Pontis *et al*., 2019, 2022). These observations led to the hypothesis that KZFPs may primarily be involved in preventing the spread of TEs, which represents a threat during the genome reprogramming that takes place in the germline and during zygotic genome activation (Hancks and Kazazian, 2016; Kim *et al*., 2016; Durnaoglu, Lee and Ahnn, 2021). While parallels in the evolutionary turnover of TEs and KZFPs lend support to this hypothesis (Jacobs *et al*., 2014), several lines of evidence suggest that the arms race model is overly simplistic. Notable findings include widespread and tightly regulated expression of KZFPs in somatic tissues, their targeting of TEs that had long lost transposition ability, signs of selection for KZFP-binding motifs on some TE subfamilies, and the enrichment of KZFP binding sites at gene promoters (Imbeault, Helleboid and Trono, 2017; de Tribolet-Hardy *et al*., 2023). These led to the proposal that KZFPs are domesticators of the regulatory potential of TEs for the benefit of the host (Friedli and Trono, 2015; Trono, 2015). In line with this model, we demonstrated that the KZFP/TE system regulates gene expression in adult tissues (Ecco *et al*., 2016), and that KZFPs partner up with their TE targets to shape the evolution of gene regulatory networks (Imbeault, Helleboid and Trono, 2017; Helleboid *et al*., 2019). Further support for broad roles for KZFPs in regulating gene expression came from the finding that about 20% human family members contain variant KRAB sequences unable to recruit TRIM28 (Helleboid *et al*., 2019; Tycko *et al*., 2020; Stoll *et al*., 2022; Milovanović *et al*., 2026), and that some of them display distinct functionalities resulting from the presence of additional domains such as SCAN and DUF3669 (Rosspopoff, Trono and Feschotte, 2025).

Recent phylogenetic studies indicate that at least 25% cis-regulatory elements (CREs) and transcription factor (TF) binding sites in the human genome reside in TEs, a majority of them of recent evolutionary origin (Du *et al*., 2024). In addition, some KZFPs bind directly to gene promoters, without identifiable underlying TE-derived sequence (Imbeault, Helleboid and Trono, 2017; Yang *et al*., 2017; Farmiloe *et al*., 2020, 2023; de Tribolet-Hardy *et al*., 2023; Forey *et al*., 2025; Pulver *et al*., 2025). While it may occasionally reflect the progressive erasure of TE signature motifs in these regions (Matsushima, Planet and Trono, 2024), comparative analyses indicate that some promoter-proximal KZFP binding sites exhibit accelerated sequence evolution, implying that these promoters are subjected to selective pressures to integrate KZFP-regulated networks (Farmiloe *et al*., 2023).

Recent studies have assigned specific functions to a handful of KZFPs, from early development and imprinting to organogenesis and the control of stress responses. (Yang *et al*., 2017; Pontis *et al*., 2019; Forey *et al*., 2025; Fueyo *et al*., 2025; S. Mesquita *et al*., 2025 as examples). These results already underline a key principle: KZFPs do not exhibit significant functional redundancy, which is also supported by systematic differences in their genomic targets, expression patterns, and domain composition. Thus, each of the ∼380 human KZFPs should be regarded as a potentially distinct contributor to human biology warranting focused investigation.

As a step towards this long-term endeavour, we sought to establish a scalable approach to assign functional roles to individual KZFPs. We reasoned that a standardized perturbation-based approach could provide an entry point to identify KZFPs with measurable biological impact and generate hypotheses regarding their mechanisms of action. In this study we performed an arrayed screen of 366 human KZFPs, assessing the consequences of their inducible overexpression on the growth and viability of human erythroleukemia K562 cells. We selected cell proliferation as a quantitative and integrative readout of cell-state perturbation, reasoning that ectopic expression of KZFPs capable of substantially reshaping transcriptional programs would produce detectable effects on cellular fitness. This strategy enabled us to compare systematically family members within a uniform experimental context and to identify candidates exerting the strongest phenotypic effects. We then combined transcriptomic profiling, chromatin occupancy analyses, data mining, and proteomic approaches to characterize a subset of these candidates. Together, this work establishes a structured framework to extract key functional insights from individual KZFPs. We anticipate that this approach will facilitate the rational exploration of this large and diverse transcription factor family and guide more focused investigations in individually relevant biological contexts.

## Results

### A high-throughput screen to identify human KZFPs affecting cell proliferation

To assess the impact of KZFP expression on cellular proliferation, we individually transduced human erythroleukemia K562 cells with doxycycline-inducible lentiviral vectors encoding 366 human KZFPs. Following puromycin selection, we cultured transduced cells in the presence or absence of doxycycline and monitored cell populations at 4-, 7-, and 9-days post-induction using PrestoBlue, which measures metabolic activity as a surrogate for cell viability and proliferation (Fig. 1A, S1A, S1B). Overexpression of approximately 25% of the tested KZFPs (89/366) induced a >15% reduction in PrestoBlue signal at day 9, indicating impaired cell growth (Fig. 1B, 1C, S1C). Notably, these KZFPs did not share any obvious common feature. They spanned a broad range of evolutionary ages (Fig. 1D), showed no binding bias toward promoters or TEs (Fig. 1E), and their binding sites were not enriched for any particular TE families (Fig. 1F). However, a modest correlation was noted between the number of transcription start sites (TSS) bound by KZFPs and the drop in Presto blue signal induced by their overexpression (Fig. 1G), and SCAN-containing KZFPs (SKZFPs) tended to induce proliferation defects more frequently than family members lacking this domain (Fig. 1H).

**Figure 1:**
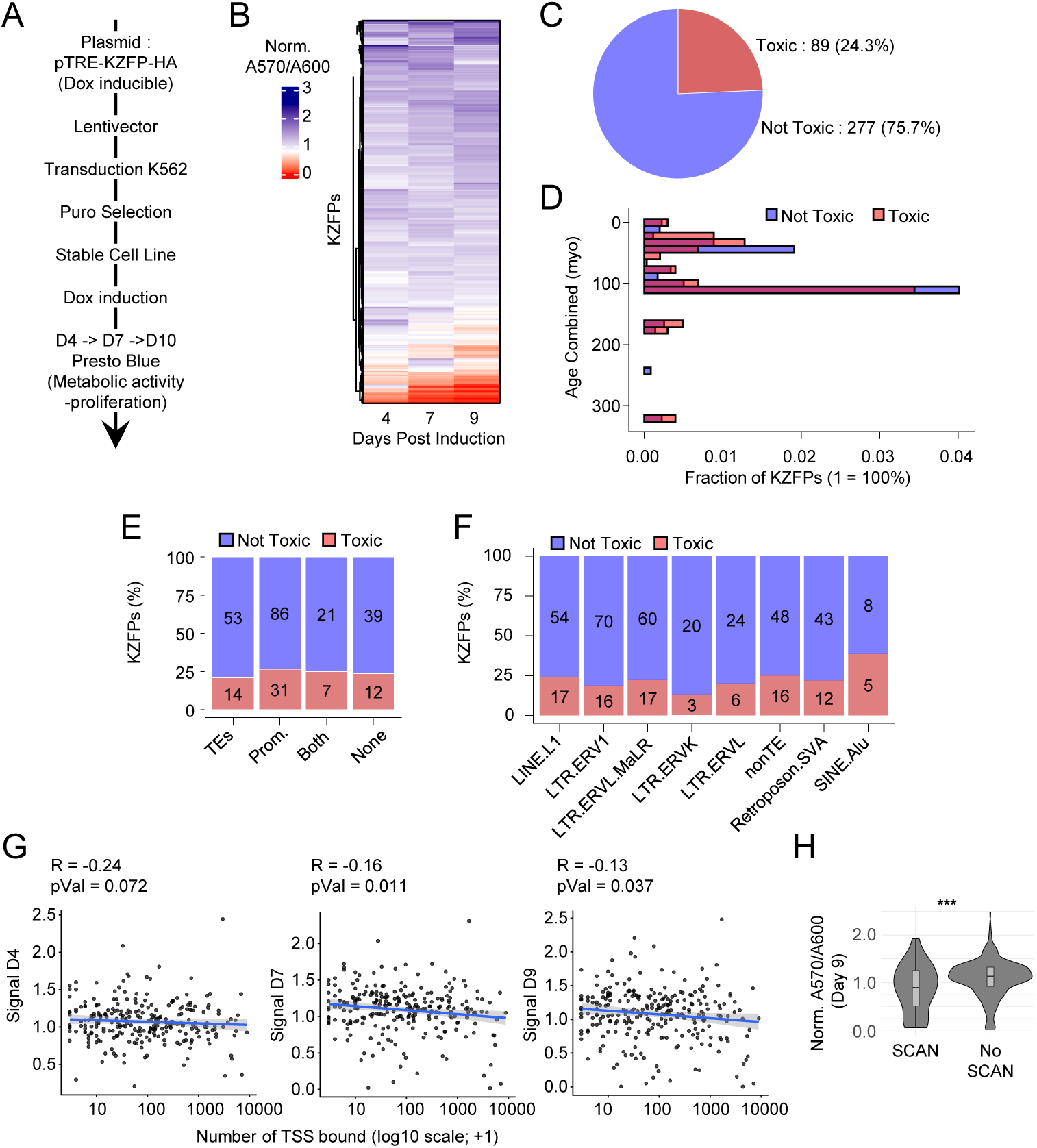
A high-throughput screen to identify human KZFPs interfering with cell proliferation. (A) Overview of the procedure for establishing the effect of KZFPs overexpression (OE) over proliferation. K562 cells were transduced, followed by puro selection, Doxycycline induced activation and measurement of proliferation after 4, 7 and 9 days of OE. The proliferation is assessed with presto blue treatment followed by absorbance measurement. (B) Heatmap representing the normalized absorbance along the timepoints. Raw values A570/A600A (background-corrected) were normalized to the mean of GFP/LacZ controls per batch, then to the corresponding ±Dox condition at each timepoint. (C) percentage and absolute number of KZFPs classified after normalization using a threshold of 0.85 (15% decrease). Red indicates toxic KZFPs (≤ 0.85), and blue indicates non-toxic KZFPs (> 0.85). (D) Distribution of evolutionary ages for KZFPs classified as toxic (red) and non-toxic (blue). (E) Percentage of toxic, (red) and non-toxic, (blue) KZFPs binding to promoters or TEs. Absolute numbers depicted on the graph for each category. (F) Percentage of toxic, (red) and non-toxic, (blue) KZFPs binding to major TE families. Absolute numbers depicted on the graph for each category. (G) Normalized absorbance for K562 overexpressing KZFPs relative to number of TSSs bound. (H) Normalized absorbance for K562 overexpressing SCAN containing KZFPs (SKZFPs), or other KZFPs for 9 days.

Following these observations, we focused subsequent analyses on a subset of four KZFPs with distinct evolutionary ages, genomic targets, and domain architectures. This group comprises three promoter-enriched KZFPs (ZNF257, ZNF498/ZSCAN25, and ZNF18), and the TE-binding ZNF43 (Fig. S1E). They range in evolutionary age from ∼180 million years (ZNF18) to ∼29 million years (ZNF257), spanning early mammalian to primate evolution. (Fig S1F). Among these four KZFPs, ZNF43 and ZNF257 harbour a canonical TRIM28-recruiting KRAB domain consistent with transcriptional repression, whereas ZNF18 and ZNF498 contain a variant KRAB domain together with an upstream SCAN domain (https://krabopedia.org/kzfp/).

### LTR/ERV1-repressing ZNF43 regulates genes involved in fatty acid metabolism and detoxification

ZNF43 is a ∼43-million-year-old KZFP with a canonical TRIM28-recruiting KRAB domain and 19 zinc fingers that preferentially recognize an *LTR/ERV1*-embedded sequence (Fig. S1D). We first verified that ZNF43 overexpression impaired the growth of K562 cells (Fig. S2A, B) and profiled their transcriptome by deep RNA sequencing (RNA-seq). To identify potential direct targets of ZNF43, we selected genes that were downregulated upon ZNF43 overexpression (Adjusted p-value < 0.05) and harboured a ZNF43 binding site within 10 kb of their TSS (Fig. 1A). Eight genes met these criteria, each displaying a nearby ZNF43-binding *LTR/ERV1* integrant: *DNAI4, ECI1, GSTO1, APOL1/APOL2* (genes in tandem with a ZNF43 binding site in between)*, HYAL1, NFU1* and *CLTB* (Fig. 2B, S2C, D). *DNAI4* encodes for a component of the axoneme (Zhang *et al*., 2019). In contrast, the other seven targets of ZNF43-mediated repression encode proteins linked to fatty acid metabolism and detoxification: ECI1 supports lipid β-oxidation (van Weeghel *et al*., 2012), GSTO1 maintains redox balance (Board, 2011), APOL1 and APOL2 regulate lipid transport and immunity (Pays, 2021), HYAL1 remodels the extracellular matrix (Garantziotis and Savani, 2019; Lu et al., 2025), NFU1 assembles mitochondrial iron–sulfur clusters (Navarro-Sastre et al., 2011) and CTLB is a constituent of endocytic vesicles (Kirchhausen *et al*., 1987). We then used the ENCODE database (https://www.encodeproject.org/) (Kagda *et al*., 2025) to characterize the chromatin state at these eight loci in K562 cells, which express low baseline levels of ZNF43. This analysis revealed enrichment of TRIM28 and H3K9me3 at three ZNF43 binding sites (near *DNAI4*, *GSTO1* and *APOL1/2*), contrasting the predominance of active marks (H3K27ac, H3K4me3, H3K4me1, H3K9ac and H3K36me3) over the others (Fig. S2E). To explore the physiological contexts in which ZNF43 functions, we analyzed tissue-specific transcriptomes from the Human Protein Atlas database (proteinatlas.org) (Karlsson *et al*., 2021). *ZNF43* is highly expressed in thymus, bone marrow and ovary but silenced in liver (Fig. 2C, D, S2F). Expression of five of the eight ZNF43 target genes show the opposite pattern, with high transcript levels in the liver (Fig. 2C, D, S2F), an organ that plays a central role in detoxification and fatty acid metabolism. Consistent with this expression pattern, the KZFP/KAP1-associated SETDB1-mediated H3K9me3 repressive mark was found to be enriched in thymus and depleted in liver at LTR/ERV1 integrants located close to the *ECI1*, *GSTO1*, *APOL1/2*, *HYAL1*, *NFU1* and *CLTB* TSS (Fig. 2E, F, S2G). We conclude from these data that ZNF43 regulates a transcriptional program related to fatty acid metabolism and detoxification, allowing for the preferential expression of its effectors in the liver (Fig. 2G). Interestingly, neither expression nor chromatin state followed the same pattern at the functionally unrelated *DNAI4* locus, indicating that this gene is subjected to other dominant regulators.

**Figure 2:**
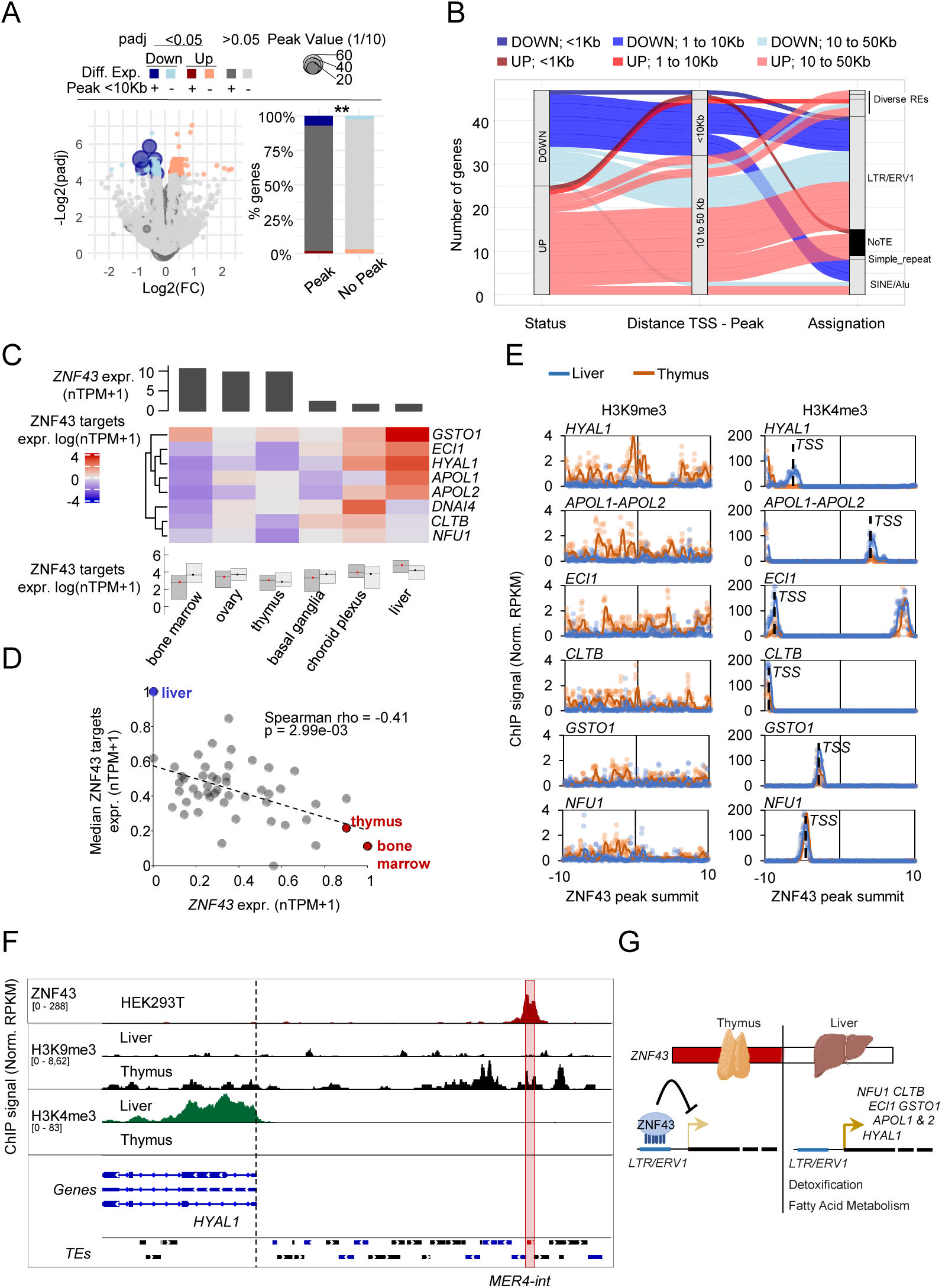
A tissue-specific regulation of fatty acid and detoxification genes via ZNF43 and its targeted TEs. (A) Gene expression changes upon OE of HA-tagged ZNF43 in K562 cells. Volcano plots represent log fold change *vs* -Log10(Adj.pVal). Differentially expressed genes (pAdj ≤ 0.05) are coloured (blue: down; red: up). Dark points denote genes with a proximal ZNF43 peak (MACS2; ± 10 kb around TSS). Dot size indicates peak score. Proportions of genes in each category are displayed. Percentage of genes assigned to each category are represented beside the associated volcano plots (Fisher’s exact test). (B) ZNF43 peaks classification related to associated differentially expressed (DE) genes based on 1/ fold change (blue = down, red = up), 2/ distance between the peak and the TSS (≤ 1Kb, 1Kb≤ d≤ 10Kb, 10Kb≤ d≤50Kb) and 3/ targets (Repetive Elements, family of transposable Elements, Other). (C) **Top**: Bar plot showing *ZNF43* expression across six human tissues representing the most extreme *ZNF43* expression levels (three highest and three lowest)(Karlsson *et al*., 2021); Protein Atlas, proteinatlas.org). **Middle**: Heatmap of ZNF43 target gene expression (log (nTPM + 1)) across the same six tissues. **Bottom**: Distribution and median expression of ZNF43 target genes (dark grey) compared to randomized gene sets (light grey) across these tissues. (D) *ZNF43* expression *vs* ZNF43 targets averaged expression across human tissues. Spearman correlation was calculated (rho = -0.41; Adj.pVal ≤0.05). (E) H3K9me3 and H3K4me3 in liver and thymus https://www.encodeproject.org/ (Kagda *et al*., 2025) ±10Kb around ZNF43 peak submit *TSS* of ZNF43 targets are indicated. (F) Integrated Genome Viewer (IGV) screenshots showing the H3K9me3 and H3K4me3 in liver and thymus of two individual subject https://www.encodeproject.org/ (Kagda *et al*., 2025) at the *HYAL1* TSSs. Position of the ZNF43 peak is indicated as well as the *MER4-int* integrant providing the binding site. (G) A model for tissue-specific regulation of fatty acid and detoxification genes via ZNF43 and its targeted TEs

### Intronic binding motif propagation enables ZNF257 to regulate a recently evolved MAGEA gene cluster

ZNF257 is a ∼29-million-year-old canonical KZFP with 11 zinc fingers and is enriched at gene promoters (Fig. S1E). When overexpressed, it markedly interfered with the proliferation of K562 cells (Fig. S3A, B). RNA-seq profiling performed 72 hours after its induction revealed that genes harbouring a ZNF257 peak within 1 kb of their TSS were significantly more frequently downregulated than others, indicating that ZNF257 functions as a promoter-proximal repressor (Fig. 3A, B). Consistent with other promoter-enriched KZFPs, ZNF257-bound regions display significantly lower PhyloP conservation scores than surrounding sequences, indicating accelerated sequence turnover at promoters of genes that predate the emergence of ZNF257 (Farmiloe *et al*., 2023). This supports the proposal that ZNF257 responsiveness was acquired through rapid diversification of promoter sequences. To explore ZNF257 activity in its physiological context, we analysed single-cell transcriptomic datasets from the Human Protein Atlas project, which revealed that *ZNF257* expression is most prominent in testis-associated germ cell populations, particularly spermatogonia and spermatocytes (Fig. 3C, S3C). Putative ZNF257 gene targets display heterogeneous levels of expression across germ cell types (Fig. 3C, S3C). However, the *MAGEA* (Melanoma associated antigen A) gene cluster, located on the X chromosome (Fig. S3D), attracted our attention owing to i) its exclusive expression in spermatogonia and spermatocytes (Fig. 3C, D, S3C), ii) its marked downregulation in ZNF257-overexpressing K562 cells (Fig. 3E), and iii) the binding of ZNF257 at several places within its locus (Fig. 3F). *MAGE* genes probably emerged before the divergence of Protochordata from Chordata. Although a single *MAGE* is found in fish, frog and chicken, most eutherians have multiple families of MAGE genes, likely the combined result of retrotranspositions and gene duplications (Katsura and Satta, 2011). The oldest member of the *MAGEA* cluster, *MAGEA6*, emerged ∼91million years ago (Fig. 3G) yet it contains within its first intron a binding site for the much younger ZNF257, implying that it is only secondarily that it became regulated by this KZFP (Fig. 3H, S3E). The MAGEA locus then underwent extensive tandem duplications, propagating the ZNF257 binding sequence, nucleated by a conserved 5′-GAGGCA-3′ motif, to several places within the resulting gene cluster (Fig. 3I, J, S3F). Parallel analysis of *ZNF257*, *MAGEA3* and *MAGEA6* (the two most highly expressed and most ZNF257-sensitive MAGEA genes) alongside spermatozoids differentiation show that their expression peaks in spermatogonia, while that of *ZNF257* peaks at the spermatocyte stage (Fig. 3K). It strongly suggests that ZNF257 contributes to initiating the transcriptional repression of these two MAGEA genes during early spermiogenesis, after which their silencing may be stabilized through stable epigenetic mechanisms such as DNA methylation.

**Figure 3:**
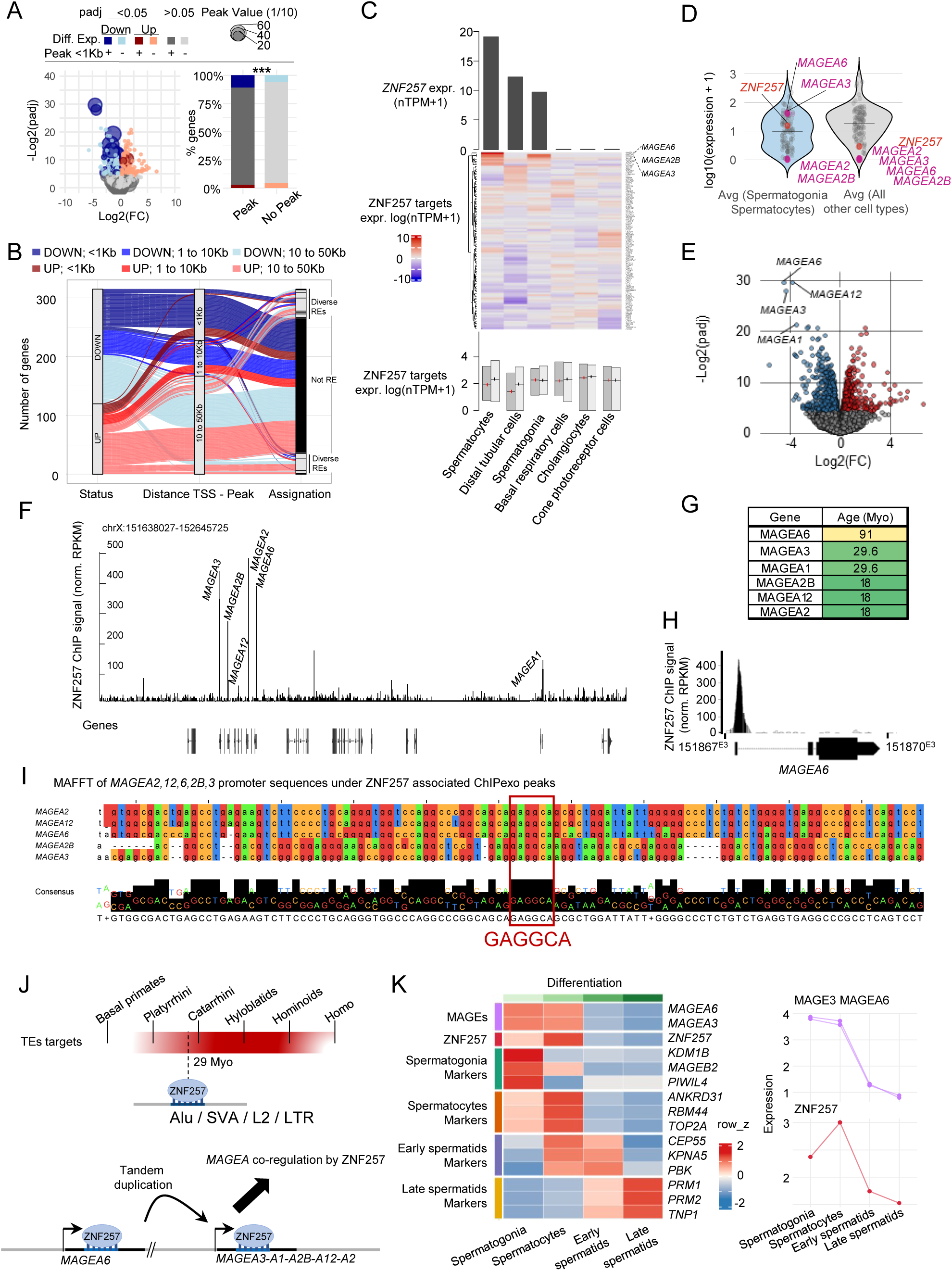
Intronic binding motif propagation enables ZNF257 to regulate a recently evolved MAGEA gene cluster. (A) Gene expression changes upon OE of HA-tagged ZNF257 in K562 cells. Volcano plots represent log fold change *vs* -Log10(Adj.pVal). Differentially expressed genes (Adj.pVal ≤ 0.05) are colored (blue: down; red: up). Dark points denote genes with a proximal ZNF257 peak (MACS2; ± 1 kb around TSS). Dot size indicates peak score. Proportions of genes in each category are displayed. Percentage of genes assigned to each category are represented beside the associated volcano plots (Fisher’s exact test). (B) ZNF257 peaks classification related to associated DE genes based on 1/ fold change (blue = down, red = up), 2/ distance between the peak and the TSS (≤1Kb, 1Kb≤d≤10Kb, 10Kb≤d≤50Kb) and 3/ targets (Repetive Elements, family of transposable Elements, Other). (C) **Top**: Bar plot showing *ZNF257* expression across six cellular subtyped representing the most extreme *ZNF257* expression levels (three highest and three lowest) (Karlsson *et al*., 2021); Protein Atlas, proteinatlas.org. **Middle**: Heatmap of ZNF257 target gene expression (log (nTPM + 1)) across the same six cellular subtyped. **Bottom**: Distribution and median expression of ZNF257 target genes (dark grey) compared to randomized gene sets (light grey) across these cellular subtyped. (D) Expression of *ZNF257* (orange dot) and ZNF257 targets defined as downregulated genes (Adj.pVal ≤0.05) harbouring a Peak at ≤ 1kb from the TSS), across spermatogonia and spermatocytes cellular subtypes (averaged), or any other cellular subtypes (averaged). *MAGEA* genes targeted by ZNF257 are highlighted in pink. (E) Gene expression changes upon OE in K562 cells for ZNF257. The volcano plots represent fold changes relative to control (x-axis) and -Log10(Adj.pVal) (y-axis). Coloured dots indicate DE genes, (blue: down; red: up). Members of the MAGEA family are labelled. (F) Integrated Genome Browser (IGB) screenshots showing the ZNF257 ChIP-exo track in 293T cells (Imbeault *et al*., 2017), at the MAGEA cluster. MAGEA genes are annotated above associated ZNF257 peaks. (G) Evolutionary age of MAGEA genes targeted by ZNF257 (H) Zoomed-in view of the ZNF257 ChIP-exo peak at the *MAGEA6* locus. (I) Multiple sequence alignment (MAFFT) of DNA sequences underlying ZNF257 ChIP-exo peaks at MAGEA genes. (J) A model for the regulation of a recently evolved MAGEA gene cluster by ZNF257 (K) Normalized RNA counts of *MAGEA6*, *MAGEA3, ZNF257* and known markers of spermatogenesis across testis cellular subtypes (Karlsson *et al*., 2021); Protein Atlas, proteinatlas.org.

### ZNF498 is a transcriptional activator that regulates microtubule cytoskeleton organization linked to neuronal function

Having observed that SCAN-containing KZFPs (SKZFPs) were prone to impair K562 cell proliferation (Fig. 1H), we explored two of them in greater depth. ZNF498 is a 105 myo SKZFP that contains a SCAN and a variant KRAB domain upstream of seven zinc fingers. ChIP-exo analysis in 293T cells indicates that ZNF498 is preferentially recruited to promoters. (Fig S1E) Its overexpression markedly impaired K562 cell growth (Fig. S4A, B), with RNA-seq performed at day 3 of doxycycline addition revealing the induction of genes harbouring a ZNF498 peak within 1 kb of their TSS (Fig. 4A, B). Thus, ZNF498 appears to act as a promoter-proximal transcriptional activator. Notably, ZNF498 is broadly expressed across multiple cell types in the central nervous system, including microglia, astrocytes, oligodendrocytes, oligodendrocyte precursor cells (OPCs), and both excitatory and inhibitory neurons (Fig. 4C, S4C). Many of its putative transcriptional targets exhibit a similar pattern, particularly in excitatory neurons where they display higher expression levels compared to randomly selected controls (Fig. 4C, S4C). Gene Ontology analysis of putative ZNF498 target genes point to regulation of microtubule cytoskeleton organization as a prominent functional theme (Fig. 4D). Key genes in this category include *LRRC4B*, *BBX*, *STMN3*, *ZNF467*, *SKA1*, *RTTN*, *DYRK1A*, *ROCK1*, and *PPP4R2*.

**Figure 4:**
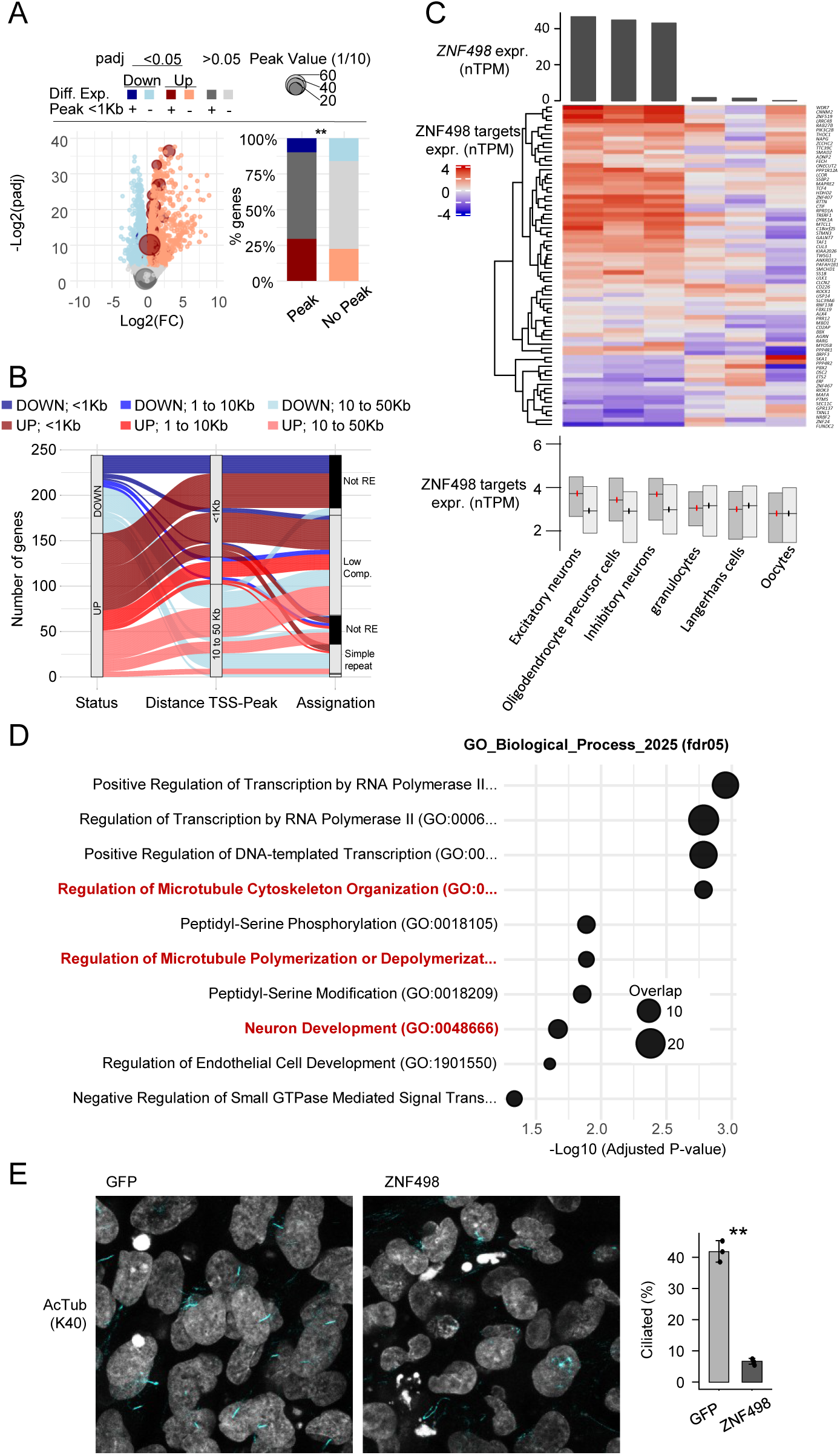
ZNF498 activates genes involved in microtubules cytoskeleton Organization. (A) Gene expression changes upon OE of HA-tagged ZNF498 in K562 cells. Volcano plots represent log fold change vs -Log10(Adj.pVal). Differentially expressed genes (Adj.pVal ≤ 0.05) are colored (blue: down; red: up). Dark points denote genes with a proximal ZNF498 peak (MACS2; ± 1 kb around TSS). Dot size indicates peak score. Proportions of genes in each category are displayed. Percentage of genes assigned to each category are represented beside the associated volcano plots (Fisher’s exact test). (B) ZNF498 peaks classification related to associated DE genes based on 1/ fold change (blue = down, red = up), 2/ distance between the peak and the TSS (≤1Kb, 1Kb≤d≤10Kb, 10Kb≤d≤50Kb) and 3/ targets (Repetive Elements, family of transposable Elements, Other). (C) **Top**: Bar plot showing *ZNF498* expression across six cellular subtyped representing the most extreme *ZNF498* expression levels (three highest and three lowest) (Karlsson *et al*., 2021); Protein Atlas, proteinatlas.org. **Middle**: Heatmap of ZNF498 target gene expression (log (nTPM + 1)) across the same six cellular subtyped. **Bottom**: Distribution and median expression of ZNF498 target genes (dark grey) compared to randomized gene sets (light grey) across these cellular subtyped. (D) Over-representation of ZNF498 targets in Gene Ontology (GO) Biological Process (BP) terms ranked (x axis) according to enrichment significance -Log10(Adj.pVal). (E) Observation of primary cilia formation defect in non-proliferating RPE1 upon ZNF498-HA OE (GFP as control). Quantification on the right (Statistical test: t.test).

Microtubule cytoskeleton organization is essential for cellular viability and the formation of specialized cellular structures, particularly in excitatory neurons, where axon maintenance depends on tightly regulated microtubule dynamics. The primary cilium is another microtubule-based organelle whose assembly relies on precise cytoskeletal organization, making it a sensitive readout for perturbations in microtubule-associated pathways.

To test functionally whether ZNF498 overexpression disrupts microtubule-dependent processes, we turned to the well-established and experimentally tractable model of primary cilium formation in hTERT-RPE1 cells. In this model, ciliogenesis can be robustly induced and quantitatively assessed in differentiated non-proliferating cells (Fig. S4D). Upon lentivector-mediated overexpression of ZNF498, primary cilium formation was severely impaired in differentiating RPE1 cells (Fig. 4E). This phenotype correlated with the upregulation of ZNF498 target genes involved in microtubule organization (*LRRC4B*, *BBX*, *STMN3* and *ZNF467)* (Fig. S4E). Together, these results identify ZNF498 as a transcriptional activator of gene modules controlling cytoskeleton-dependent and suggest that this TF may act as a regulator of neuronal cytoskeletal architecture, warranting investigation in relevant neural models.

### ZNF18 controls events important for spermatogenesis

We finally focused on ZNF18, a ∼164 myo SKZFP containing a SCAN and a variant KRAB domain, like ZNF498, followed by five zinc fingers that govern preferential association with gene promoters (Fig S1E). ZNF18 overexpression markedly impaired K562 cell proliferation (Fig. S5A, B). RNA-seq performed 72 hours after doxycycline induction revealed that, as for ZNF498, genes harbouring a ZNF18 peak within 1 kb of their TSS were preferentially upregulated (Fig. 5A, B). To define the physiological context of ZNF18 activity, we examined its expression across human cell types and found that it is prominently expressed during spermatogenesis, particularly at the early and late spermatid stages (Fig. 5C, S5C). A subset of putative ZNF18 target genes, defined as being upregulated upon overexpression and harbouring a ZNF18 peak within 1 kb of their TSS, is also highly expressed in these germ cell populations (Fig. 5C, S5C). Their functional annotation points to chromatin organization and DNA repair (*BANF1*, *SPOP*, *SWI5*), and to cytoskeletal remodelling (*DYNC2I2*, *NSL1*, *TRIM37*, *WIPF2, GRK5*) as processes regulated by ZNF18. Both chromatin remodelling and cytoskeletal reorganization are hallmarks of spermiogenesis, during which the nucleus undergoes extensive compaction and specialized microtubule-based structures such as the flagellum are assembled. The co-expression of ZNF18 and its target genes at the spermatid stage suggests that ZNF18 activates a transcriptional program supporting these processes.

**Figure 5:**
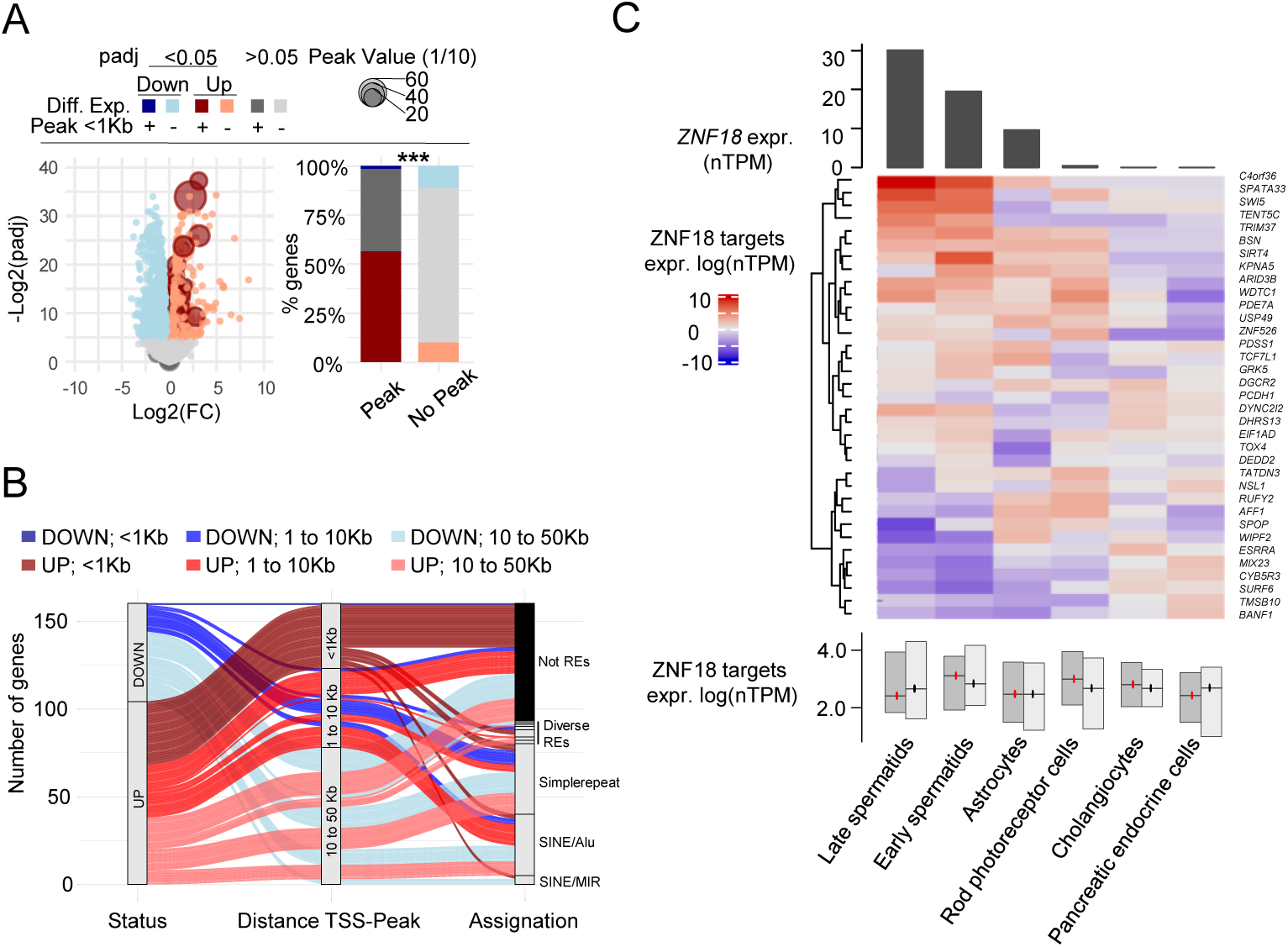
ZNF18 activates genes important for spermatogenesis. (A) Gene expression changes upon OE of HA-tagged ZNF18 in K562 cells. Volcano plots represent log fold change vs -Log10(Adj.pVal). Differentially expressed genes (Adj.pVal ≤ 0.05) are colored (blue: down; red: up). Dark points denote genes with a proximal ZNF18 peak (MACS2; ± 1 kb around TSS). Dot size indicates peak score. Proportions of genes in each category are displayed. Percentage of genes assigned to each category are represented beside the associated volcano plots (Fisher’s exact test). (B) ZNF18 peaks classification related to associated DE genes based on 1/ fold change (blue = down, red = up), 2/ distance between the peak and the TSS (≤1Kb, 1Kb≤d≤10Kb, 10Kb≤d≤50Kb) and 3/ targets (Repetive Elements, family of transposable Elements, Other). (C) **Top**: Bar plot showing *ZNF18* expression across six cellular subtyped representing the most extreme *ZNF18* expression levels (three highest and three lowest) (Karlsson *et al*., 2021); Protein Atlas, proteinatlas.org. **Middle**: Heatmap of ZNF18 target gene expression (log (nTPM + 1)) across the same six cellular subtyped. **Bottom**: Distribution and median expression of ZNF18 target genes (dark grey) compared to randomized gene sets (light grey) across these cellular subtyped.

### The SCAN domain assembles distinct transcriptional complexes

Given that both ZNF18 and ZNF498 act as transcriptional activators, we next sought to determine how their SCAN domain contributes to this non-canonical KZFP activity. The SCAN domain is a conserved ∼80-amino-acid oligomerization module found in a subset of C₂H₂ zinc finger transcription factors, including 24 human KZFPs (Matsushima *et al*., 2025). Unique to vertebrates, it mediates selective homo- and hetero-dimerization among SCAN-containing proteins *via* a helical bundle interface structurally related to a retroviral capsid (Liang *et al*., 2012; Rimsa, Eadsforth and Hunter, 2013). This dimerization capacity enables the combinatorial assembly of protein complexes, and SZFPs have accordingly been proposed to function as modulatory or scaffolding factors that integrate distinct regulatory pathways (Williams, Blacklow and Collins, 1999; Schumacher *et al*., 2000; Sander *et al*., 2003; Urrutia, 2003; Huang *et al*., 2019). Interestingly, the SCAN-encoding sequences of ZNF18 and ZNF498 have been under strong purifying selection during evolution, suggesting that they play key roles in the functions of these SKZFPs (Fig. S6A-D). In contrast, the KRAB domain of ZNF18 appears less constrained, and it is truncated in ZNF498 (Fig. S6A-D). Accordingly, deleting the SCAN domain completely abrogated or partially alleviated the proliferation defect induced by ZNF18 and ZNF498 overexpression, respectively (Fig. 6A, B). To identify the potential protein partners of ZNF498 and ZNF18, we performed immunoprecipitation followed by mass spectrometry (co-IP–MS) on their full-length and SCAN-deleted variants. In each case, we identified a set of SCAN-dependent interactors, *i.e.* proteins that co-immunoprecipitated with the full-length construct but were absent in controls and lost upon deletion of the SCAN domain (Fig. 6C). We detected 99 SCAN-dependent interactors common to ZNF18 and ZNF498, which represents a significant enrichment compared to random permutations (Fig. 6D). Amongst them, other SCAN-containing proteins were strongly overrepresented as expected (Fig. 6E-G). We also observed that deleting the SCAN domain of either SKZFP led to a consistent loss of proteins interaction involved in the regulation of gene expression, including core transcriptional machinery components and co-activators (Fig. 5H, I). Furthermore, the individual SCAN dependant interactor of these SZFPs were enriched in factors involved in DNA repair for ZNF18 (Fig. S6E-G) and in basal transcriptional machinery and Mediator complex components for ZNF498 (Fig. S6H-J).

**Figure 6:**
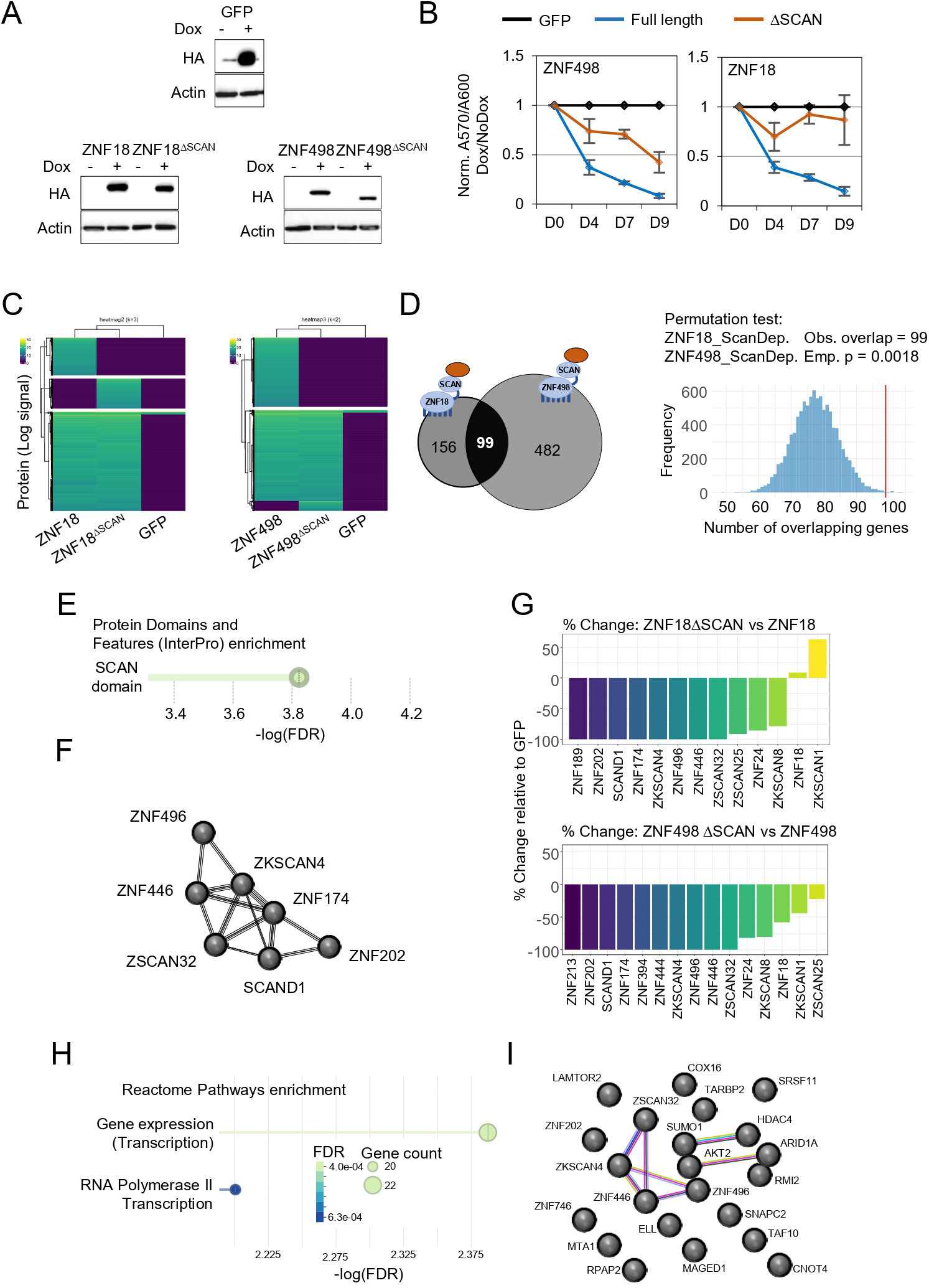
KZFPs with non-canonical modes of action. (A) HA signal after OE of HA-tagged ZNF18, ZNF18^ΔSCAN^, ZNF498, ZNF498^ΔSCAN^ or GFP in K562 cells. Actin as control. (B) Metabolic activity as a surrogate for the proliferation of K562 overexpressing HA-tagged ZNF18, ZNF18^ΔSCAN^, ZNF498, ZNF498^ΔSCAN^ or GFP for 4, 7 and 9 days using PrestoBlue (Black = GFP, Blue = Full Length, Orange = delta-SCAN). (C) Heatmap representing protein abundance detected by mass spectrometry after co-immunoprecipion of HA tagged HA-tagged ZNF18, ZNF18^ΔSCAN^, ZNF498, ZNF498^ΔSCAN^ or GFP in K562 cells. K-means clustering was performed and cluster were annotated manually to differentiate 3 interactors subtypes based on SCAN domain dependency. (D) Overlap of proteins lost upon SCAN deletion (SCAN dependant interactors) for ZNF498 and ZNF18. On the right is the result of random permutation test on SCAN dependant interactors for ZNF498 and ZNF18. Histogram represent result of 10 000 permutation. Observed overlap (99) is represented with a red line. (E) Over-representation of SCAN dependant interactors in Protein Domain and Feature (InterPro) terms ranked (x axis) according to signal. Dot size represent protein count. (F) STRING network of SCAN containing proteins interacting with both ZNF18 and ZNF498. (G) Histogram illustrating the % of change caused by SCAN ablation for ZNF18 or ZNF498. (H) Over-representation of SCAN dependant interactors in Reactome pathways terms ranked (x axis) according to FDR. Dot size represent protein count. (I) STRING network of SCAN dependant interactors with both ZNF18 and ZNF498 (add color code of the Edges).

## Discussion

This work presents a systematic functional survey of the human KZFP family, combining arrayed overexpression screening with integrative multi-omic characterization. Of 366 human KZFPs tested, approximately 25% impaired the proliferation of K562 cells. As K562 cells do not represent the endogenous context for most of these factors, the observed fitness effects likely reflect the capacity of these KZFPs to forcefully reshape transcriptional programmes, a property that, paradoxically, makes ectopic expression in a permissive but non-native model a useful discovery tool. KZFPs with sufficient regulatory potency to perturb cellular fitness outside of their normal setting are strong candidates for playing important roles within it. Detailed follow-up of four such candidates, ZNF43, ZNF257, ZNF498 and ZNF18, revealed, as hypothesised distinct modes of action, ranging from TE-linked transcriptional repression to promoter-proximal gene silencing and transcriptional activation. These findings reinforce the view that KZFPs, while often viewed as a homogeneous family of TE-repressive TFs, are rather functionally diverse regulators with wide-ranging impacts on human biology.

The four KZFPs characterised here illustrate this diversity. ZNF43 represses a coherent set of genes involved in fatty acid metabolism and detoxification through binding to nearby LTR/ERV1 integrants, with its expression anticorrelating that of its targets: i.e., highly expressed in thymus and bone marrow, where these metabolic genes are silent, and lowly expressed in liver, where they are most active. This represents a clear example of host genomes coopting TE-derived sequences and shaping their regulatory activities in a cell-type specific manner by the differential expression of KZFPs. ZNF257, by contrast, acts as a promoter-proximal repressor whose targets show accelerated sequence evolution at their promoters, consistent with integration into a KZFP-orchestrated GRN through rapid promoter diversification, a feature previously described for KZFPs (Farmiloe *et al*., 2023). Its regulation of the MAGEA gene cluster exemplifies a distinct evolutionary mechanism: an ancestral intronic binding site, present in MAGEA6 gene body, before ZNF257 emerged, was propagated across the cluster through tandem duplication, enabling coordinated regulation of multiple paralogs. Temporal expression analysis during spermatogenesis further suggests that ZNF257 initiates MAGEA repression at the spermatogonia-to-spermatocyte transition, after which silencing may be maintained through epigenetic mechanisms such as DNA methylation. ZNF498 and ZNF18, both SCAN-containing KZFPs with variant KRAB domains, on the other hand acted as transcriptional activators. ZNF498 activates a programme centred on microtubule cytoskeleton organisation, as demonstrated by the disruption of ciliogenesis upon its overexpression, and both ZNF498 and its targets are broadly expressed in the central nervous system, particularly in excitatory neurons where microtubule dynamics are essential for axonal architecture. ZNF18 similarly activates genes involved in chromatin remodelling and cytoskeletal reorganisation at the spermatid stage, processes that are hallmarks of spermiogenesis. Together, these case studies demonstrate that even within a single screen for a single phenotype, KZFPs with fundamentally different regulatory logics can be identified and then mechanistically dissected to uncover their unique properties.

The characterisation of ZNF498 and ZNF18 highlights the SCAN domain as a central determinant of their function in this setting as its deletion abolished or reduced the proliferation phenotype. Proteomics analysis revealed that the SCAN domain mediates recruitment of a shared core of interactors, including other SCAN-containing proteins and components of the transcriptional machinery and co-activator complexes, while also assembling protein-specific partners: DNA repair factors for ZNF18 and Mediator complex components for ZNF498. These findings support a model in which SCAN-containing KZFPs operate through protein assemblies distinct from the classical KRAB/TRIM28 repressive pathway, consistent with the emerging view that variant KRAB domains and accessory domains such as SCAN and DUF3669 have enabled functional diversification within the KZFP family (Helleboid *et al*., 2019; Matsushima *et al*., 2025; Rosspopoff, Trono and Feschotte, 2025). The observation that the SCAN domain can recruit transcriptional activators rather than repressors provides a molecular explanation for why the two SKZFPs studied here upregulate rather than silence their targets, and raises the broader question of how many of the 24 human SZFPs, SKZFPs and more broadly variant KRAB containing KZFPs share this activating potential.

### Limitations of the study

Several methodological considerations bear on the interpretation of these results. The use of cellular proliferation as a screening readout enabled discovery of KZFPs linked to diverse biological processes, but it is not pathway-specific and therefore provides prioritisation rather than direct functional resolution. The arrayed format also captures KZFP function in only one cellular context, and many KZFP family members likely act in specific tissues, developmental stages, or stimulus conditions not represented in K562 cells. However, comparison of KZFP and target gene expression across tissues and cell types yielded biologically coherent hypotheses that can now be tested in the appropriate physiological systems. Looking ahead, the natural extension of this work is a pooled, multi-context screening. Interrogating KZFP function across multiple cell lines, differentiation trajectories, and perturbation conditions should capture the context-dependent activities that an arrayed screen in a single cell type inevitably misses. Combined with the functional, evolutionary, and proteomic data generated here, such approaches will accelerate the systematic annotation of this vast transcription factor family. If the four KZFPs characterised here are any indication, each member of the family harbours a distinct regulatory logic waiting to be uncovered.

## Methods

### Cell culture

K562 cells were maintained in RPMI 1640 medium supplemented with 10% fetal bovine serum (FBS) and 1× penicillin–streptomycin. HEK293T cells were cultured in Dulbecco’s modified Eagle’s medium (DMEM) supplemented with 10% FBS and 1× penicillin–streptomycin. hTERT-RPE1 cells were cultured in DMEM supplemented with 10% FBS and 1× penicillin–streptomycin. All cells were maintained at 37°C in a humidified incubator with 5%.

For ciliogenesis experiments, hTERT-RPE1 cells were plated to reach high confluence before serum withdrawal. Primary cilium formation was induced by culture in low-serum medium for 48–72 h, allowing cells to exit the cell cycle and enter a non-proliferative state permissive for ciliogenesis.

### Cloning of inducible KZFP expression constructs

A collection of 366 human KZFP open reading frames was assembled for inducible overexpression screening (de Tribolet-Hardy *et al*., 2023). Coding sequences were cloned into doxycycline-inducible lentiviral transfer vectors designed to express N-terminally HA-tagged proteins. GFP and LacZ were used as controls throughout the screen and in downstream validation experiments. Selected constructs encoding ZNF43, ZNF257, ZNF498 and ZNF18 were used for follow-up mechanistic studies. For SCAN-domain functional analyses, deletion constructs lacking the SCAN domain were generated for ZNF18 and ZNF498 in the same lentiviral backbone. All constructs were verified by Sanger sequencing before use.

### Lentiviral production and transduction

Lentiviral particles were produced in HEK293T cells by transient co-transfection of transfer, packaging and envelope plasmids. Cells were transfected at approximately 70–80% confluence using a standard lipid-based transfection reagent. Viral supernatants were collected 48 h after transfection, cleared by centrifugation, filtered through 0.22-µm membranes, and used fresh or stored appropriately until use. Recipient K562 or hTERT-RPE1 cells were transduced under conditions optimized for efficient gene delivery. Transduced cells were selected with puromycin for several days, after which doxycycline was added to induce expression of the HA-tagged transgene.

### Arrayed overexpression screen

To systematically assess the effect of human KZFP overexpression on cellular fitness, K562 cells were individually transduced with doxycycline-inducible lentiviral vectors encoding 366 human KZFPs. The screen was performed in an arrayed 96-well format, with each KZFP assayed in technical triplicate under induced (+Dox) and non-induced (−Dox) conditions. GFP and LacZ control wells were included on each plate to account for plate-to-plate and batch-to-batch variability. Peripheral wells were filled with medium to reduce evaporation-driven edge effects.

Following puromycin selection, doxycycline was added to induce expression of the HA-tagged KZFPs. Proliferation was assessed after 4, 7 and 9 days of induction. At each time point, metabolic activity was measured using PrestoBlue^TM^ reagent according to the manufacturer’s instructions. Absorbance was recorded at 570 nm and 600 nm using a plate reader, and the A570/A600 ratio was used as a surrogate for viable cell number and proliferative capacity.

For computational normalization, raw A570/A600 values were background-corrected, then normalized in two steps. First, each value was divided by the mean of GFP and LacZ controls from the same batch and induction condition. Second, the resulting value was divided by the corresponding −Dox condition for the same KZFP and time point, yielding a relative proliferation score. KZFPs with a normalized proliferation score ≤ 0.85 at day 9 were classified as proliferation-impairing within this screening framework.

### Validation of selected KZFPs by proliferation assay

ZNF43, ZNF257, ZNF498 and ZNF18 were selected for follow-up validation based on their growth-inhibitory phenotype, genomic binding profiles, evolutionary ages and domain architecture.

Independent K562 populations expressing the corresponding HA-tagged proteins were generated by lentiviral transduction and puromycin selection. Expression was induced with doxycycline, and proliferation was monitored at days 4, 7 and 9 using the same PrestoBlue-based assay and normalization framework as in the primary screen. GFP-expressing cells served as controls.

For SCAN-domain analyses, K562 cells expressing HA-tagged ZNF18, ZNF18^ΔSCAN^, ZNF498, ZNF498^ΔSCAN^ or GFP were cultured in parallel, and relative proliferation was quantified as above.

### Western blotting

Cells were lysed in RIPA buffer supplemented with protease inhibitors. Lysates were clarified by centrifugation at high speed at 4°C, and protein concentration was determined using a standard colorimetric or fluorometric assay. Equal amounts of protein were mixed with denaturing loading buffer, heated, separated by SDS–PAGE, and transferred to nitrocellulose membranes.

Membranes were blocked in 5% milk in TBST and incubated with Anti-HA-Peroxidase (High Affinity, 50 mU/mL, Roche, ref: 12013819001) or horseradish peroxidase (HRP)-conjugated anti-actin antibody (1:5,000 dilution, ThermoFisher, ref: MA5-15739-HRP). After three washes, protein bands were visualized using enhanced chemiluminescence detection reagent (Advansta Inc, ref: K-12049-D50) and imaged using a chemiluminescent imaging system (Fusion FX from Vilber).

### RNA extraction and RNA-seq library preparation

For transcriptome profiling, K562 cells expressing the indicated inducible constructs were treated with doxycycline for 72 h before harvest. Total RNA was extracted using the NucleoSpin RNA plus kit (Macherey-Nagel) according to the manufacturer’s recommendations. RNA quantity and purity were assessed by spectrophotometry, and RNA integrity was evaluated before library preparation.

RNA-seq libraries were prepared using the Illumina TruSeq Stranded mRNA kit. Libraries were sequenced in 75 bp paired-end formats on the NovaSeq 6000 sequencers.

### RNA-seq processing and differential expression analysis

RNA-seq libraries were prepared using the Illumina TruSeq Stranded mRNA kit. Libraries were sequenced in 75 or 100 bp paired-end formats on the Illumina HiSeq 4000 and NovaSeq 6000 sequencers, respectively. RNA-seq reads were mapped to the hg19 human genome releases using hisat v2.1.060. Only uniquely mapped reads were used for counting over genes and repetitive sequence integrants (MAPQ > 10). Counts for genes and TEs were generated using featureCounts v2 and normalized for sequencing depth using the TMM method implemented in the limma package of Bioconductor. Counts on genes were used as library size to correct both gene and TE expression. For repetitive DNA elements, an in-house curated version of the Repbase database was used. Differential gene expression analysis was performed using Voom61 as implemented in the Limma package of Bioconductor62. Genes with an adjusted P value ≤ 0.05 were considered significantly differentially expressed. For downstream figure generation, differential expression outputs were integrated with processed genomic binding tables. Volcano plots were generated in R by plotting fold-change metrics against transformed adjusted P values. Differentially expressed genes were classified as UP, DOWN or UNCHANGED according to fold-change sign and adjusted significance threshold. For ZNF43 analyses, genes with a ZNF43 peak within ±10 kb of the transcription start site were classified as peak-proximal. For ZNF257, ZNF498 and ZNF18, promoter-proximal analyses used a ±1 kb window around the transcription start site, in accordance with the definitions used in the figure panels.

Differential expression results were integrated with genomic binding profiles through a reproducible interval-processing workflow implemented with bash, awk, bedtools, and R. First, genes were annotated as differentially expressed using an adjusted P value threshold of 0.05. Differentially expressed genes were then classified as UP or DOWN according to fold-change direction.

KZFP peak files were intersected with RepeatMasker-derived repeat annotations using bedtools intersect -loj, thereby retaining all peaks while appending repeat annotation fields where overlap existed. Annotated peaks were then assigned to the closest coding-gene transcription start site using bedtools closest -d. Finally, differential expression tables and peak/TSS/repeat tables were merged on Ensembl gene identifiers to produce integrated tables for downstream figure generation.

This workflow yielded unified tables in which each gene was associated with its expression status, the nearest KZFP peak, peak score, distance from the nearest peak to the TSS, and overlap status with repetitive elements and TE families. These merged tables were used as the basis for the volcano plots, alluvial peak-classification plots, distance analyses, and TE-family analyses in Figs. 2–5 and their supplementary panels.

### Definition of putative direct KZFP target genes

To infer putative direct KZFP targets, differential expression results were integrated with previously established or reprocessed genomic binding profiles. For ZNF43, candidate direct targets were defined as genes significantly downregulated upon overexpression and associated with a ZNF43 peak within 10 kb of the transcription start site. For ZNF257, ZNF498 and ZNF18, candidate direct targets were defined as differentially expressed genes associated with a peak within 1 kb of the TSS.

Genes were additionally categorized according to the direction of expression change, distance from the nearest peak, and whether the corresponding peak overlapped repetitive DNA, a TE family annotation, or non-repetitive sequence. Distance categories used in the corresponding figure panels were ≤1 kb, 1–10 kb, 10–50 kb, and >50 kb.

### TE enrichment analysis

Transposable element annotations were obtained from RepeatMasker-derived tracks. Overlaps between KZFP peaks and TE families were computed in (de Tribolet-Hardy *et al*., 2023). For family-level enrichment analyses, the observed frequency of overlap with each TE family was compared with a background expectation using Fisher’s exact tests.

### Chromatin profiling using public datasets

To characterize the chromatin environment of KZFP-bound loci, public ChIP-seq signal tracks were retrieved from https://www.encodeproject.org/ (Kagda *et al*., 2025), (de Tribolet-Hardy *et al*., 2023) and (Brocks *et al*., 2017). Signal was visualized with deepTools using locus-centered heatmaps and average profiles.

In detail: 1/For tissue-specific ZNF43 analyses, liver and thymus H3K9me3 and H3K4me3 bigWig files were downloaded from https://www.encodeproject.org/ (Kagda et al., 2025) and relabeled locally for plotting. The following ENCODE accessions were used:

ENCFF252UZI: liver H3K9me3

ENCFF932THN: liver H3K4me3

ENCFF497GHE: thymus H3K9me3

ENCFF625VPD: thymus H3K4me3

Signal around ZNF43-centered regions was processed with computeMatrix reference-point and visualized with plotHeatmap and plotProfile from deepTools. Depending on the panel, regions were centered on ZNF43 peak summits and quantified in ±3 kb or ±10 kb windows. In the final heatmap workflow, the following parameters were used: --referencePoint center, -b 10000, -a 10000, --binSize 50, --skipZeros, and --missingDataAsZero. Heatmaps were rendered with the viridis color map and panel-specific display limits.

2/For multi-mark chromatin heatmaps centered on ZNF43-associated regions, public bigWig files (Karlsson *et al*., 2021)https://www.encodeproject.org/ (Kagda *et al*., 2025), (de Tribolet-Hardy *et al*., 2023) and (Brocks *et al*., 2017) corresponding to ZNF43, KAP1, SETDB1, H3K9me3, H3K27me3, H3K4me3, H3K4me1, H3K9ac, H3K27ac and H3K36me3 were quantified with deepTools using computeMatrix reference-point in a ±10 kb window with 50-bp bins. The workflow included --skipZeros, --missingDataAsZero, and --maxThreshold 500. Heatmaps were visualized with plotHeatmap, and sorted-region files were exported for inspection.

### Tissue-wide and single-cell expression analyses

Expression profiles of KZFPs and candidate target genes across human tissues and cell types were analyzed using Human Protein Atlas-derived tissue-consensus and single-cell expression tables (Protein Atlas, proteinatlas.org (Karlsson *et al*., 2021)). For tissue-level analyses, normalized transcript abundance values were transformed as log(nTPM + 1) for visualization. For single-cell analyses, cell subtype-resolved values were used to define the physiological contexts in which KZFP activity and target-gene repression or activation might occur.

For each selected KZFP, either all tissues/cell types or the three highest and three lowest expressing categories were displayed, depending on the figure. In the composite figure scripts, the top panel showed KZFP expression across tissues or cell types, the middle panel showed the expression of candidate target genes, and the bottom panel compared the distribution of target-gene expression to matched randomized gene sets of equal size. Randomization used fixed seeds for reproducibility. Correlations between KZFP expression and average target-gene expression across tissues were calculated using Spearman correlation for ZNF43.

### Analysis of the MAGE locus and ZNF257 occupancy

To examine ZNF257 regulation of the MAGE gene cluster, ZNF257 ChIP signal was visualized across MAGE loci using deepTools. A BED file containing MAGE-family loci was analyzed with computeMatrix scale-regions using ZNF257_pubM.bw as the signal track. The matrix was computed with 1 kb upstream and downstream flanks and a 1 kb scaled region body using the parameters --beforeRegionStartLength 1000, --afterRegionStartLength 1000, --regionBodyLength 1000, --skipZeros, --missingDataAsZero, and --sortUsing mean. Heatmaps were rendered with plotHeatmap using the viridis color map, descending region sorting, and fixed display limits (--zMin 20 --zMax 100). To assess sequence conservation and duplication-associated propagation of ZNF257-bound elements within the MAGEA cluster, promoter sequences of MAGEA genes were aligned with MAFFT using the parameters --reorder --kimura 20 --op 1.0 --maxiterate 2 --retree 1 --globalpair.

### Evolutionary analyses

Evolutionary ages assigned to KZFPs were integrated from a curated metadata table and used to compare the age distribution of proliferation-impairing and non-impairing KZFPs in the screen and to position selected candidates within the evolutionary history of the family. Additional analyses of evolutionary constraint for ZNF18 and ZNF498 used domain-wise or per-residue dN/dS estimates to compare the SCAN, KRAB and zinc-finger regions as done in (Matsushima *et al*., 2025). Functional domains were annotated based on NCBI.

### Gene ontology, pathway and domain enrichment analyses

Functional enrichment analyses were performed on gene sets derived from RNA-seq/ChIP integration. Over-representation was tested against Gene Ontology Biological Process.

### Ciliogenesis assay

To test whether ZNF498 overexpression perturbs microtubule-dependent cellular processes, hTERT-RPE1 cells were transduced with doxycycline-inducible lentiviral vectors expressing HA-tagged ZNF498 or GFP. After puromycin selection, transgene expression was induced with doxycycline and cells were serum-starved to promote ciliogenesis. Cells were fixed and stained with antibodies against ciliary markers (ARL13B), and nuclei were counterstained with DAPI. Images were acquired by fluorescence microscopy under identical conditions across samples. The fraction of ciliated cells was quantified across multiple fields and biological replicates. Differences between conditions were evaluated statistically by t-test.

### RT–qPCR

For validation of selected transcriptional changes in hTERT-RPE1 cells, total RNA was extracted following overexpression of ZNF498 or GFP control. Reverse transcription was performed using standard cDNA synthesis procedures. Quantitative PCR was performed using SYBR Green or probe-based chemistry on a real-time thermocycler. Relative transcript abundance was calculated by comparative Ct analysis after normalization to housekeeping genes.

### Affinity purification–mass spectrometry

To define the interactomes of ZNF18 and ZNF498 and assess the functional contribution of the SCAN domain, K562 cells expressing HA-tagged full-length proteins, HA-tagged ΔSCAN variants, or GFP control were induced with doxycycline and harvested for affinity purification. Lysates were prepared in immunoprecipitation buffer supplemented with protease inhibitors, clarified, and incubated with anti-HA magnetic beads overnight at 4°C. Beads were washed extensively, and bound proteins were digested on-bead using LysC and trypsin. Peptides were reduced, alkylated, acidified, desalted by StageTips or equivalent cleanup procedures, dried, and analyzed by nanoLC–MS/MS. Protein identification and quantification were performed against the human proteome using an appropriate DIA-compatible software pipeline.

### AP-MS computational analysis

Protein-group intensity matrices were processed in R from report.pg_matrix.tsv. For figure-oriented analyses, the relevant bait columns were selected according to the experimental comparison. Numeric conversion was applied to all intensity values, and missing values were replaced with 1 before log₂ transformation, matching the processing used in the figure-generation scripts. Proteins lacking signal above background were excluded and proteins were additionally required to show stronger signal in at least one bait condition than in GFP controls based on heatmap clustering (see script).

Heatmaps were generated using ComplexHeatmap, and row-wise k-means clustering was used to define interaction classes. During exploratory analysis, several values of k were evaluated, and cluster tables were exported for manual inspection. Final cluster labels were then collapsed into SCAN-dependent, other and undefined interaction classes.

Overlap between ZNF18 and ZNF498 SCAN-dependent interactors was quantified directly and statistically evaluated by Fisher’s exact tests and by random permutation. Empirical null distributions were generated from 10,000 random draws matched to the sizes of the compared sets. Overlap visualization included Venn/Euler diagrams, UpSet plots, pairwise association heatmaps and histogram-based permutation displays. Shared and factor-specific interactors were further analyzed by Reactome and InterPro enrichment and by STRING-based network visualization.

### Protein interaction network analysis

Protein–protein interaction networks were generated from interactor lists using the STRING database (https://string-db.org/ version 12.0). Interactions were filtered using a minimum confidence score cutoff of 0.4 (medium confidence). Evidence channels retained for network construction included experimental data, curated databases, co-expression, text mining, co-occurrence, gene fusion, and neighborhood evidence, as implemented in STRING. Networks were annotated based on known functional modules. Subnetwork structure was further resolved using density-based clustering (DBSCAN), enabling the identification of functionally coherent interaction modules. Functional annotation of clusters was performed using STRING-derived enrichment terms and protein annotation datasets.

### Statistical analyses

Most analyses and figure generation were performed in R; additional image-based statistics and figure assembly were performed as appropriate. Fisher’s exact test was used for enrichment analyses involving genomic feature overlaps, TE overlaps, gene-category comparisons and protein-set overlaps. Wilcoxon rank-sum tests were used for selected non-parametric group comparisons. Spearman correlation was used for tissue-wide and cell-type expression correlations. Student’s t-tests or exact t-tests were used for selected chromatin or imaging quantifications were indicated in the figure panels. For mass-spec overlap analyses, empirical P values were also estimated by permutation using 10,000 random draws. Multiple-testing correction was performed using the Benjamini–Hochberg method where appropriate. Unless otherwise stated, adjusted P values ≤ 0.05 were considered significant.

### Data visualization and software

Plots were generated using R packages including ggplot2, ggrepel, ggpubr, ComplexHeatmap, circlize, viridis/viridisLite, gridExtra, UpSetR, ggvenn, eulerr, igraph, clusterProfiler, ReactomePA, and related dependencies. Genomic interval analyses used bedtools. Chromatin heatmaps and aggregated signal profiles were generated with deepTools, including computeMatrix, plotHeatmap, and plotProfile. Genome-browser representations were prepared in IGV or IGB. Sequence alignments were performed with MAFFT.

### Data availability

Previously published ChIP-exo datasets, ENCODE chromatin datasets and Human Protein Atlas transcriptomic datasets used in this study are publicly available from their original repositories.

Genomic binding profiles of human KZFPs were obtained from previously published ChIP-exo datasets generated in HEK293T cells. Peak BED files used in this study were derived from datasets reported by Imbeault et al. and de Tribolet-Hardy et al., including resources documented through Krabopedia, except for ZNF257. For ZNF257, the original peak calling was considered suboptimal for downstream analyses and peak calling was re-computed; the re-called peak set was used for all ZNF257 analyses in the present study.

For KZFP-wide analyses, processed peak enrichment tables were used to classify KZFPs according to promoter enrichment, TE enrichment, both, or neither. Enrichment significance was assessed using Fisher’s exact tests, and multiple-testing correction was applied where appropriate.

## Code availability

Custom scripts used for screen normalization, RNA-seq/ChIP-exo integration, interval annotation, deepTools workflows, tissue-expression analyses, TE-family analyses, enrichment analyses, and AP-MS overlap analyses should be deposited in a public repository together with the processed input tables required to reproduce the published figures.

## Acknowledgements

We thank the Trono lab members for constructive discussions. Most computational work was conducted on the high-performance computing cluster developed and maintained by Scientific IT and Application Support (SCITAS) at EPFL. We thank the Gene Expression Core Facility (GECF), Flow Cytometry Core Facility (FCCF), and Proteomics Core Facility (PCF) at EPFL for technical support.

## Funding

European Research Council No. 268721 (DT)

European Research Council No. 694658 (DT)

Swiss National Science Foundation 310030_152879 (DT)

Swiss National Science Foundation 310030B_173337 (DT)

European Molecular Biology Organization (EMBO) Postdoctoral Fellowship ALTF1287-2020 (WM) Japan Society for the Promotion of Science (JSPS) Overseas Research Fellowship no. 202360326 (WM)

## Author contributions

RF. and DT. designed the research plan. RF., CR and JA designed and conducted all wet lab experiments, with technical help from SO. RF designed and oversaw the statistical analyses with input by EP. WM produced dN/dS score with input by EP. EP, pre-processed the raw NGS data. RF analyzed the data and designed the figures. RF, DT and AD wrote the manuscript, with substantial contributions from OB. and suggestions by all authors.

---

**Figure S1 – Related to Fig. 1.**
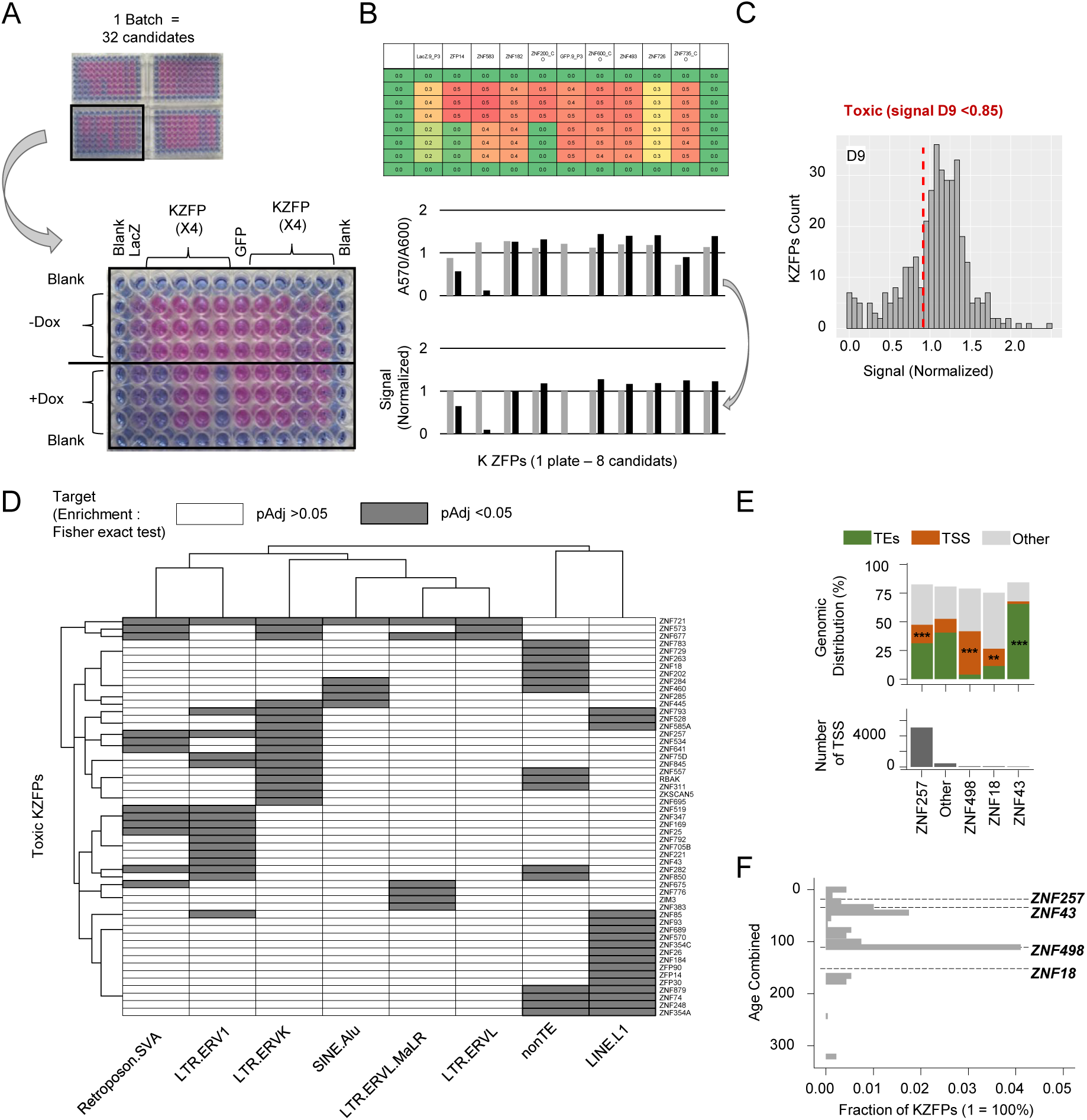
(A) Experimental workflow for one screening batch (four 96-well plates), and representative zoom of one 96-well plate after PrestoBlue coloration. Each KZFP was tested in technical triplicate under ±Dox conditions to evaluate proliferation upon induction of the HA-tagged protein. LacZ and GFP controls were included on each plate to correct for plate and batch effects. Peripheral wells were filled with PBS to minimize evaporation-driven edge effects. PrestoBlue reduced reagent appears pink in metabolically active wells and remains blue in low-viability conditions. Absorbance was measured at 570 nm and 600 nm with plate reader where 570 nm measures PrestoBlue signal and 600 nm serves as a reference wavelength to correct for background. (B) *A*570/*A*600 values were double-normalized: first, each sample was divided by the mean of all GFP and LacZ controls from the same batch (same ±Dox condition) to minimize inter-plate variability; second, values were normalized to the corresponding -Dox condition at each timepoint. This yields a timepoint-specific normalized ratio reflecting relative viability changes across conditions. (C) Distribution of normalized proliferation scores for all KZFPs screened. An arbitrary cutoff of normalized signal ≤ 0.85 was used to define “toxic” KZFPs. (D) TE family enrichment for toxic KZFPs (p ≤ 0.05). TE family enrichment assessed by Fisher’s exact test. (E) Enrichment of ChIP-exo peaks (hg19) from (Imbeault, Helleboid and Trono, 2017) and (de Tribolet-Hardy *et al*., 2023) across transposable elements (TEs) and transcription start sites (TSS) for ZNF43, ZNF257, ZNF18, and ZNF498, assessed by Fisher’s exact test. Asterisks indicate significant enrichment (*** adjusted p ≤ 0.05). The lower panel shows the absolute number of TSS bound by each factor. Other KZFPs are included as references. (F) Distribution of evolutionary ages for KZFPs. Selected candidates are highlighted.

**Figure S2 – Related to Fig. 2.**
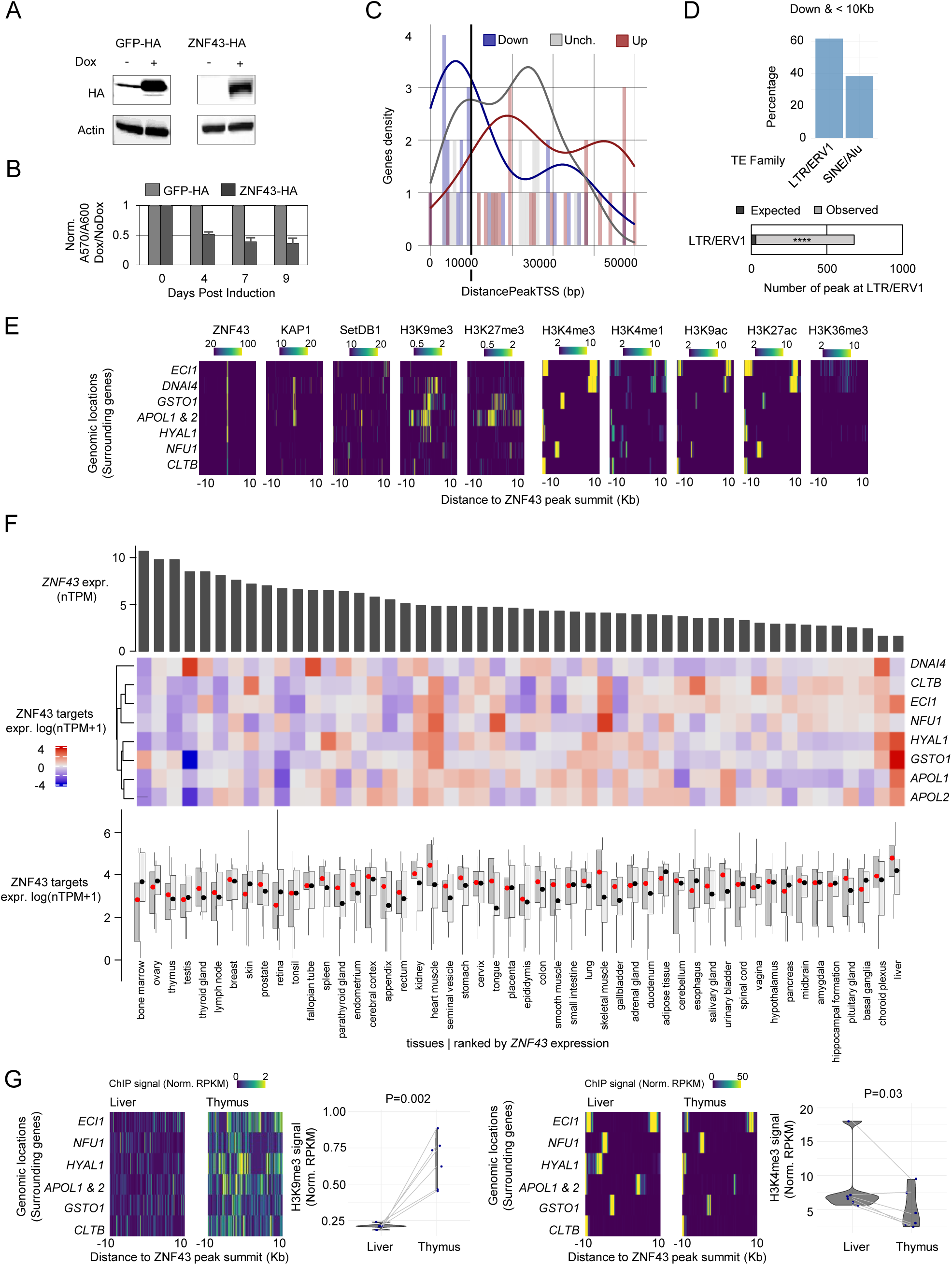
(A) Immunoblot analysis of HA-tagged GFP or ZNF43 after induction in K562 cells after 3 days. Actin used as loading control. (B) Effects of the OE of HA-tagged GFP or ZNF43 on cell proliferation after 4, 7 and 9 days. (C) Distances between DE genes (up- or downregulated; p ≤ 0.05) or non-DE genes (p > 0.05) after 3 days of ZNF43 OE and the closest ZNF43 ChIP-exo peak from HEK293T cells (Imbeault, Helleboid and Trono, 2017). (D) **Up**: TE families bound by ZNF43 within 10 kb of significantly downregulated genes (Non-exclusive – A peak of ZNF43 can overlap several TEs). **Down**: Genome-wide TE families significantly bound by ZNF43 (FDR ≤ 0.05). Absolute number of LTR/ERV1 bound by ZNF43. Overrepresentation significance calculated by Fisher’s exact test (de Tribolet-Hardy *et al*., 2023). (E) ZNF43-HA ChIPexo signal upon OE in HEK293T cells (Imbeault, Helleboid and Trono, 2017) and KAP1, SETDB1, and histone chromatin marks ChIP-seq signal in K562 cells across ZNF43 peaks associated with DE genes https://www.encodeproject.org/ (Kagda *et al*., 2025). (F) **Top**: Bar plot showing *ZNF43* expression across human tissues (Karlsson *et al*., 2021); Protein Atlas, proteinatlas.org). **Middle**: Heatmap of ZNF43 target gene expression (log (nTPM + 1)) across tissues. **Bottom**: Distribution and median expression of ZNF43 target genes (dark grey) compared to randomized gene sets (light grey) across tissues. (G) H3K9me3 and H3K4me3 in liver and thymus, https://www.encodeproject.org/ (Kagda *et al*., 2025), across ZNF43 peaks associated with DE genes. Quantification of the signal (count +/- 10Kb around ZNF43 peak submit) is presented on the right (Statistical test: Exact t.test).

**Figure S3 – Related to Fig. 3.**
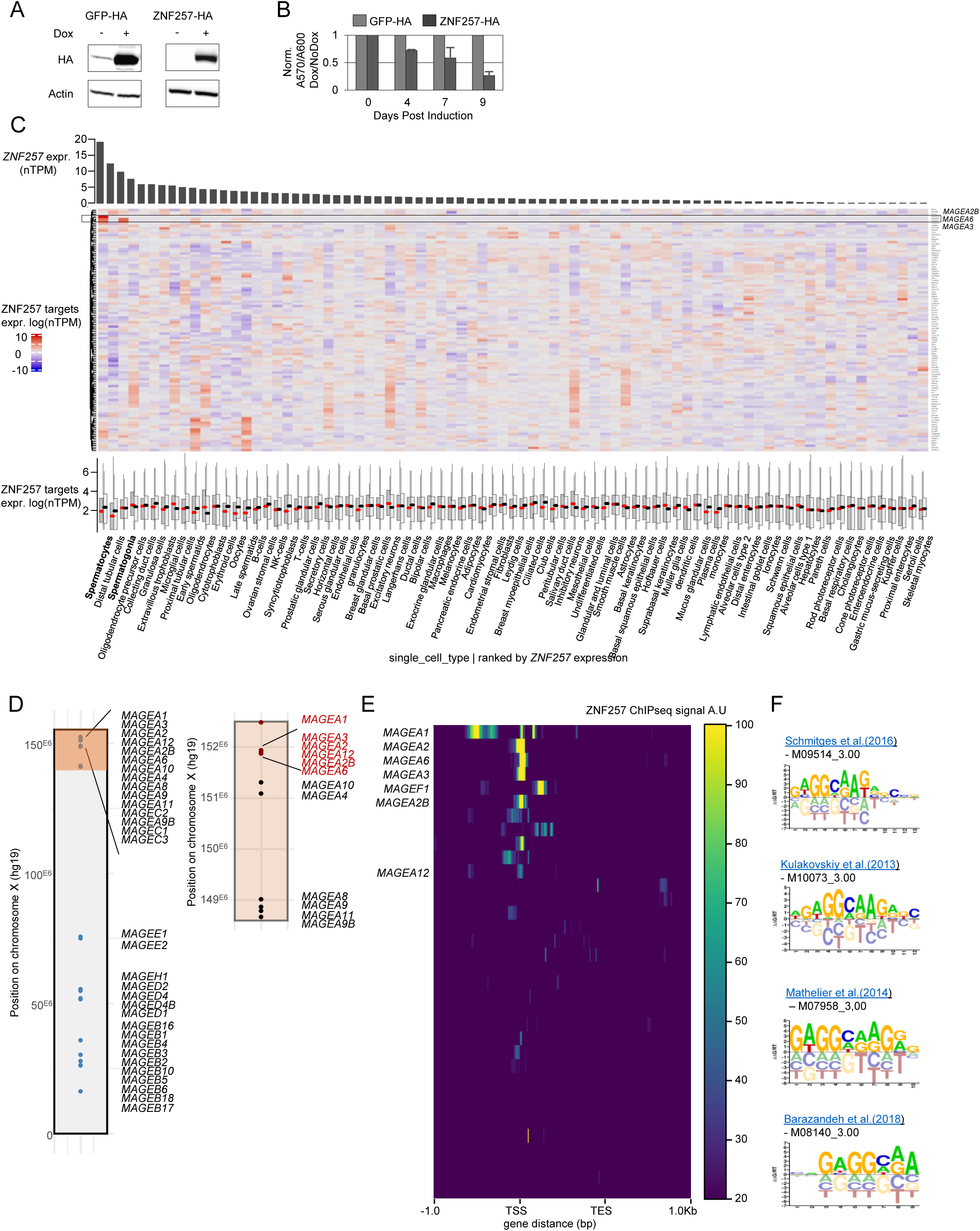
(A) Immunoblot analysis of HA-tagged GFP or ZNF257 after induction in K562 cells after 3 days. Actin used as loading control. (B) Effects of the OE of HA-tagged GFP or ZNF257 on cell proliferation after 4, 7 and 9 days. (C) **Top**: Bar plot showing *ZNF257* expression across cellular subtypes (Karlsson *et al*., 2021); Protein Atlas, proteinatlas.org). **Middle**: Heatmap of ZNF257 target gene expression (log (nTPM + 1)) across cellular subtypes. **Bottom**: Distribution and median expression of ZNF257 target genes (dark grey) compared to randomized gene sets (light grey) across cellular subtypes. (D) Genomic distribution of MAGE-encoding genes along chromosome X. The positions of MAGEA gene clusters are highlighted on the right. MAGEA genes bound by ZNF257 and downregulated following ZNF257 induction for three days are shown in red. (E) ZNF257 ChIP-exo signal at MAGEA gene. MAGEA genes targeted by ZNF257 are annotated. (F) ZNF257 DNA-binding motif reported in the CIS-BP database.

**Figure S4 – Related to Fig. 4.**
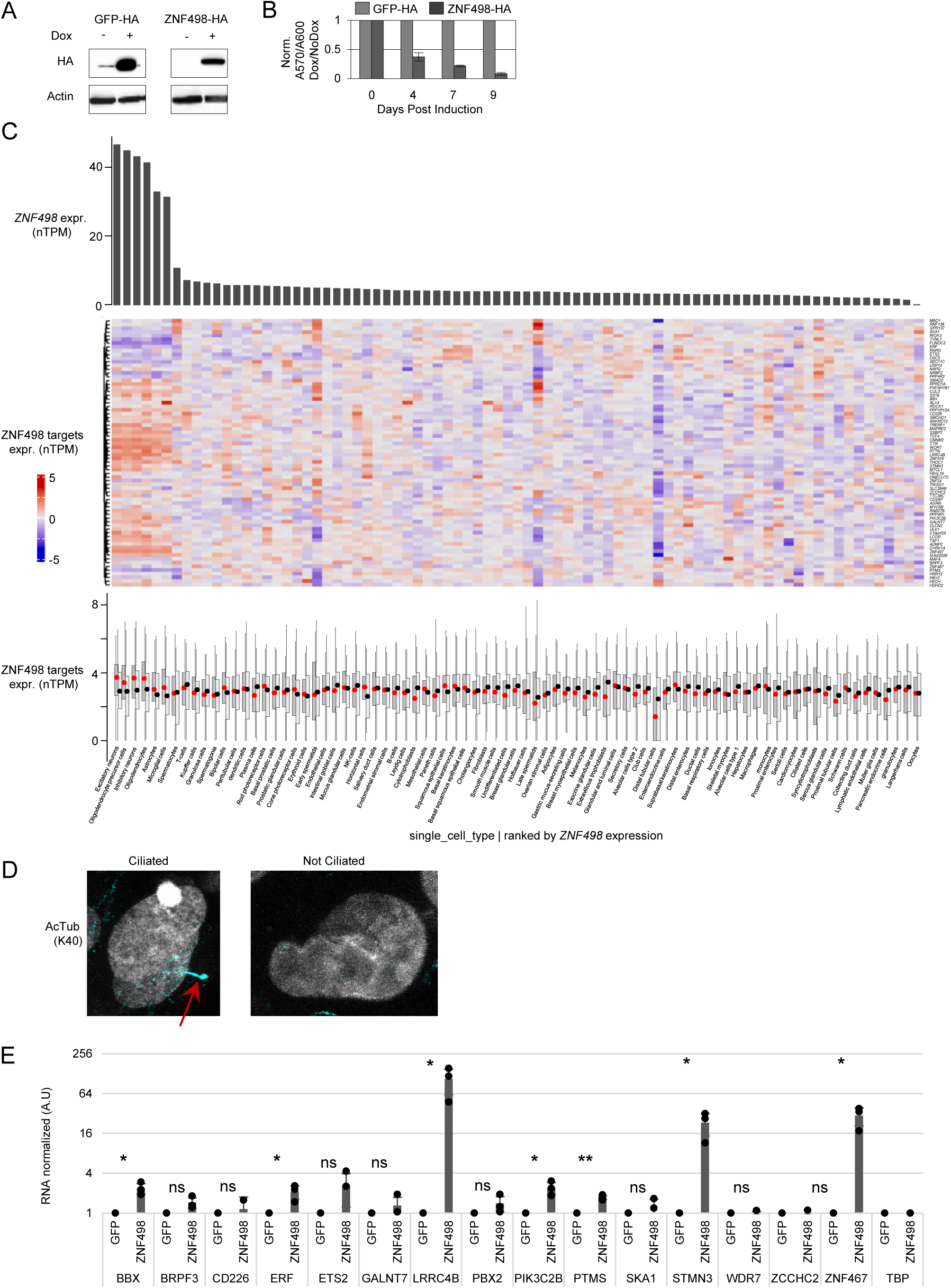
(A) Immunoblot analysis of HA-tagged GFP or ZNF498 after induction in K562 cells after 3 days. Actin used as loading control. (B) Effects of the OE of HA-tagged GFP or ZNF498 on cell proliferation after 4, 7 and 9 days. (C) **Top**: Bar plot showing *ZNF498* expression across cellular subtypes (Karlsson *et al*., 2021); Protein Atlas, proteinatlas.org). **Middle**: Heatmap of ZNF498 target gene expression (log (nTPM + 1)) across cellular subtypes. **Bottom**: Distribution and median expression of ZNF498 target genes (dark grey) compared to randomized gene sets (light grey) across cellular subtypes. (D) Example of differentiated and not proliferating RPE1 cells ciliated (left) or not (right). (E) qPCR of a subset of previously identified ZNF498 targets after OE of HA-tagged ZNF498, or GFP in RPE1 cells. (Statistical test: t.test).

**Figure S5 – Related to Fig. 5.**
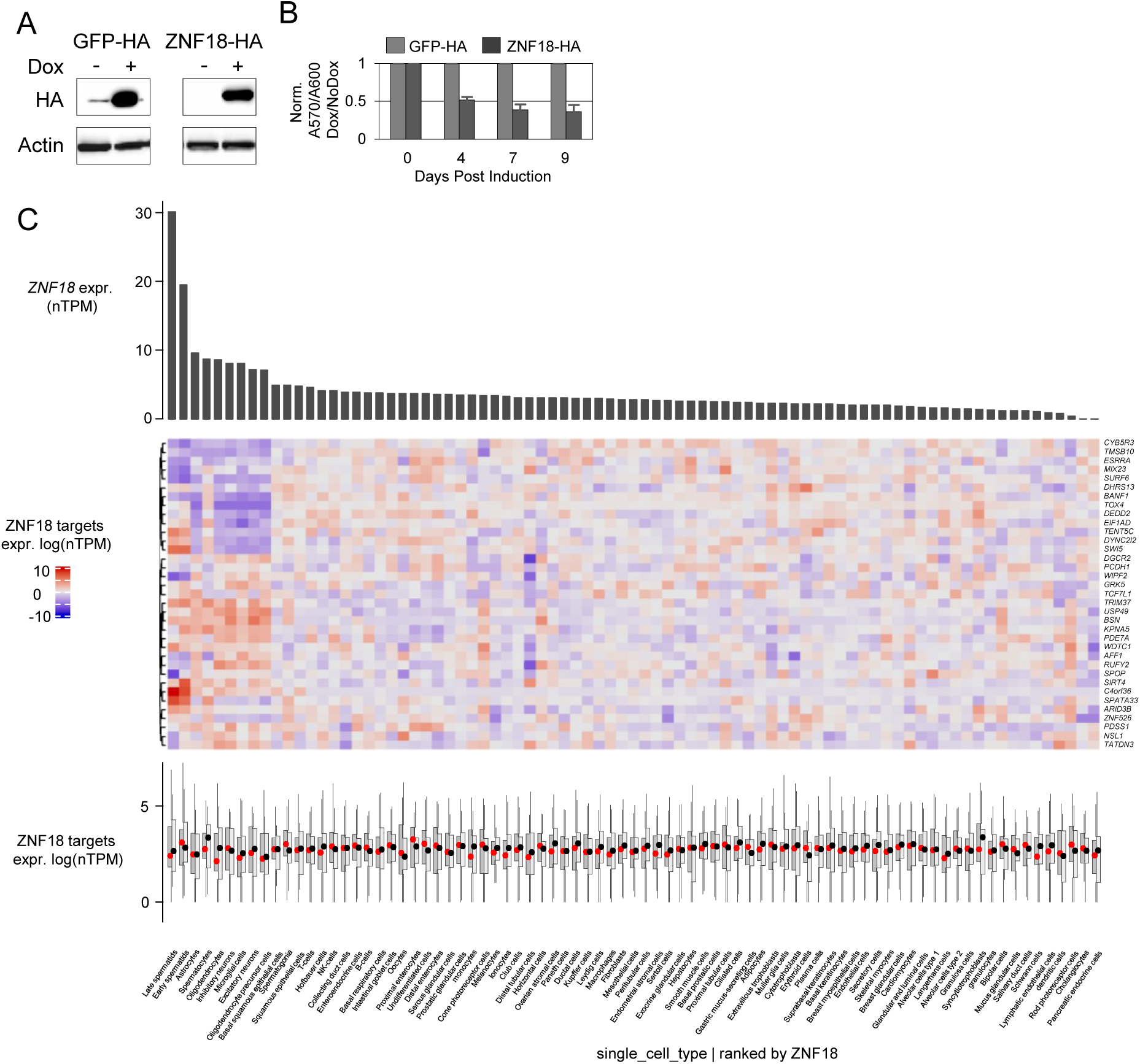
(A) Immunoblot analysis of HA-tagged GFP or ZNF18 after induction in K562 cells after 3 days. Actin used as loading control. (B) Effects of the OE of HA-tagged GFP or ZNF18 on cell proliferation after 4, 7 and 9 days. (C) **Top**: Bar plot showing *ZNF18* expression across cellular subtypes (Karlsson *et al*., 2021); Protein Atlas, proteinatlas.org). **Middle**: Heatmap of ZNF18 target gene expression (log (nTPM + 1)) across cellular subtypes. **Bottom**: Distribution and median expression of ZNF18 target genes (dark grey) compared to randomized gene sets (light grey) across cellular subtypes.

**Figure S6: Related to figure 6.**
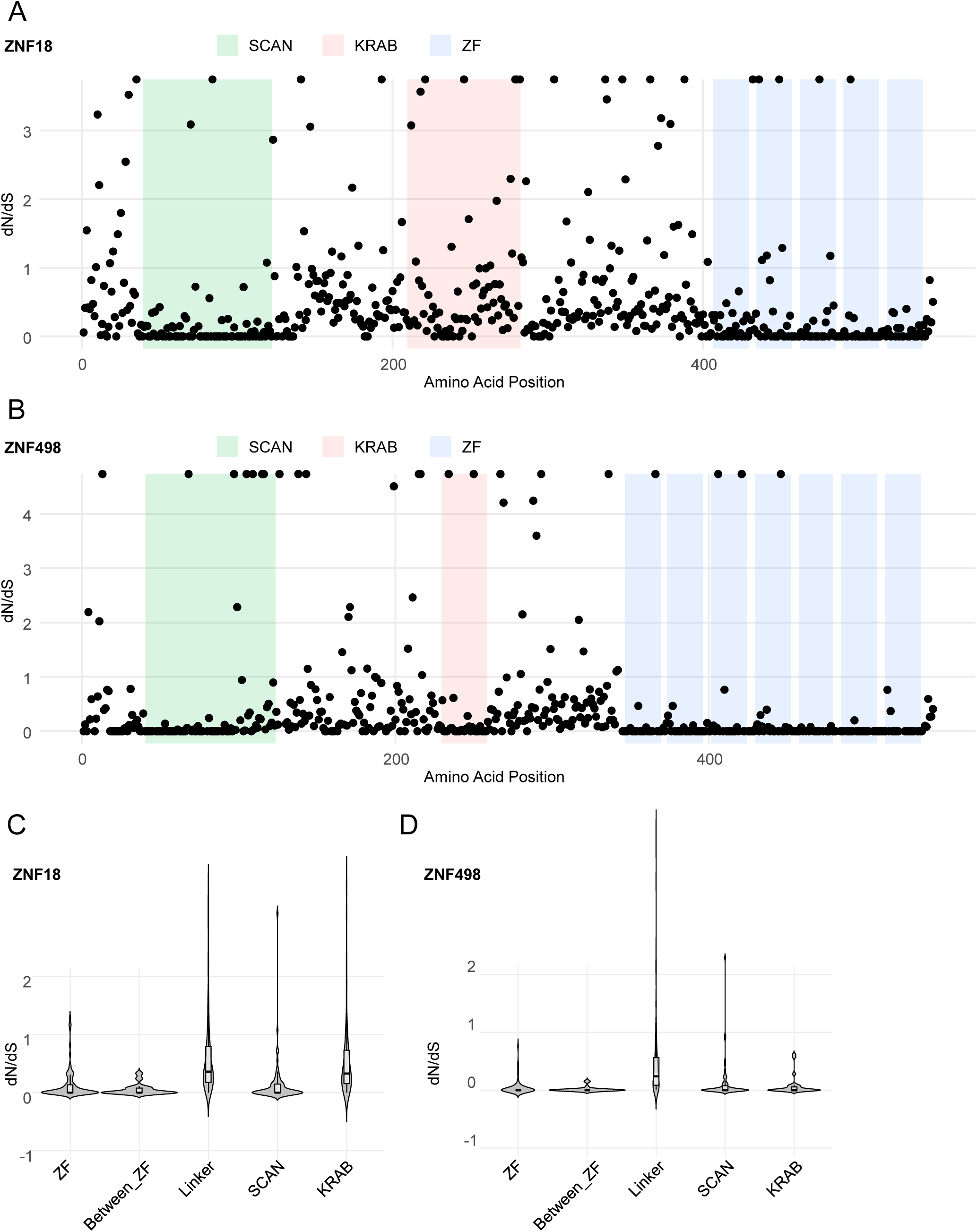
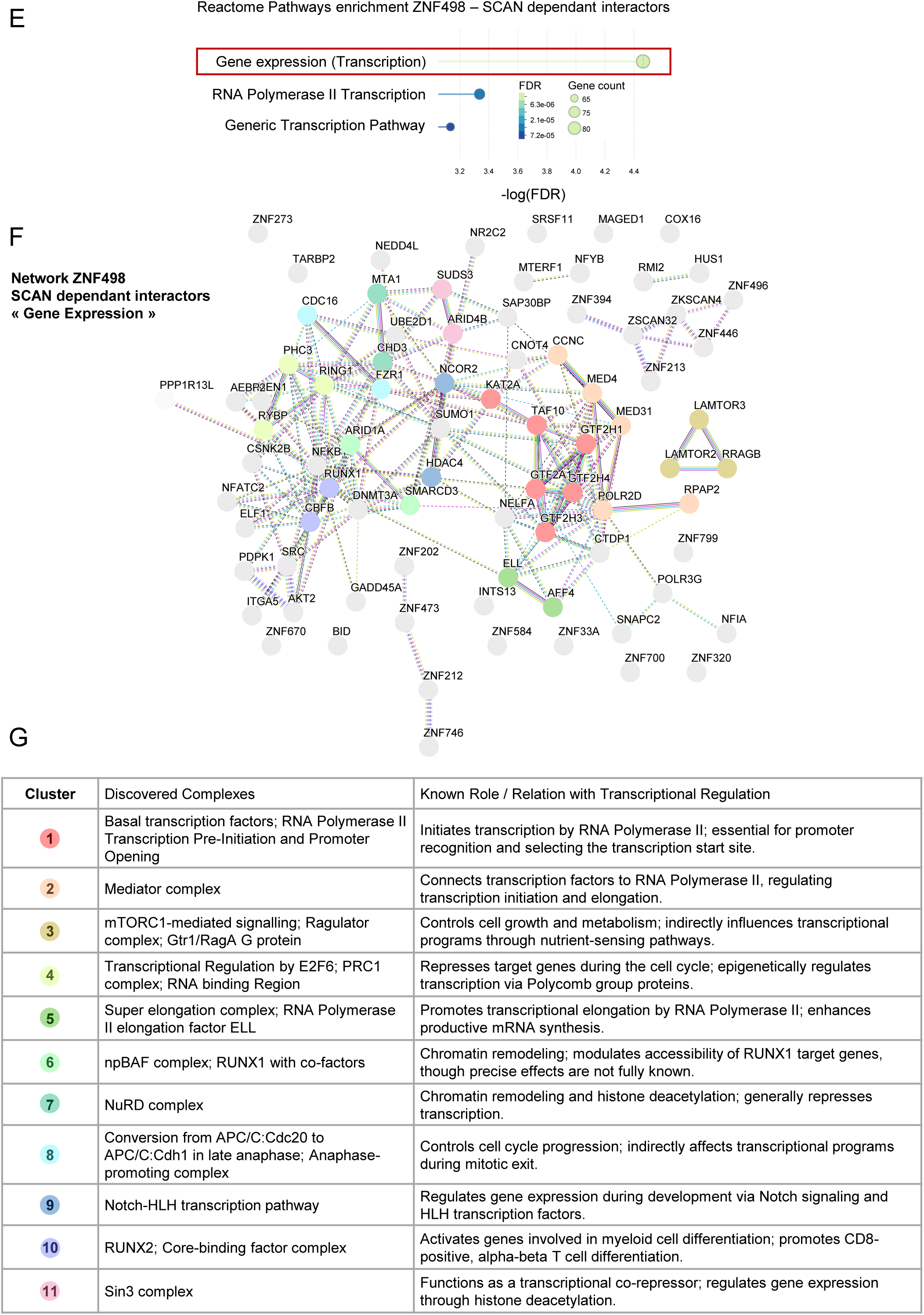
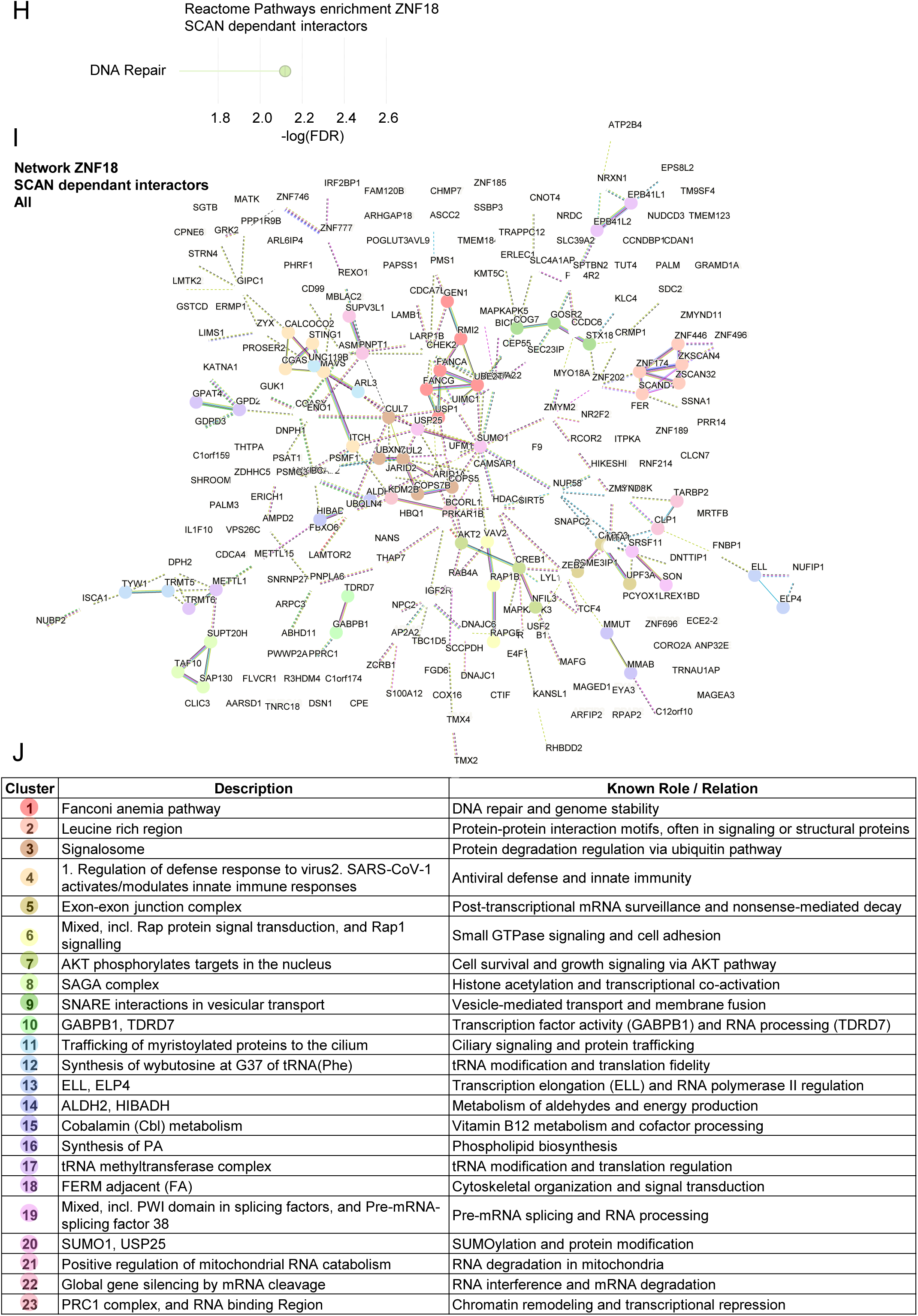
(A) dN/dS ratios per amino acid along ZNF18. Annotated functional domains: SCAN (green), KRAB (red), ZNF (blue). (B) Same as (A) for ZNF498. (C) Distribution of domain-specific dN/dS values for ZNF18 (box = IQR, bar = median). (D) Same domain-wise comparison for ZNF498. (E) Reactome pathway enrichment for ZNF498 interactors. Bubble size indicates protein count. (F) STRING network of proteins annotated in Reactome “Gene Expression.” Clustered subnetworks analysed for GO Biological Process enrichment (FDR ≤ 0.05). (G) Table reporting clustered subnetworks descriptions, and known biological roles for S6F. (H) Reactome pathway enrichment of ZNF18 interactors. Bubble size indicates protein count. (I) STRING network built using the full set of ZNF18 interactors. (J) Table reporting clustered subnetworks descriptions, and known biological roles for S6I.

